# Dysregulated Cholesterol Metabolism with Anomalous PI3K/Akt/mTOR pathway Predicts Poor Carboplatin Response in High Grade Serous Ovarian Cancer

**DOI:** 10.1101/2024.08.17.608375

**Authors:** Elizabeth Mahapatra, Arka Saha, Niraj Nag, Animesh Gope, Debanjan Thakur, Manisha Vernekar, Jayanta Chakrabarti, Mukta Basu, Amit Pal, Sanghamitra Sengupta, Sutapa Mukherjee

## Abstract

Rapidly escalating High-Grade Serous Ovarian Cancer (HGSOC) incidences, relapse, and mortalities result from failed carboplatin therapy. In this regard, reprogrammed cholesterol metabolism arising from deregulated PI3K/Akt/mTOR signaling aggravates HGSOCs to evade carboplatin. Therefore, we designed a pilot study to ascertain their clinical relevance in determining the carboplatin response of HGSOC tumors.

Non-NACT HGSOC (n=31) subjects were classified into optimum, borderline, and high cohorts based on blood cholesterol levels which positively correlated with their relative tissue cholesterol content. TCGA database showed that mutations in specific PI3K/Akt/mTOR candidates including cholesterol metabolism regulators (SREBP1, SREBP2, SRB-1, STAR, HMGCR) and prosurvival effectors (Akt, mTOR, p70S6K, P38MAPK, HIF-1α, COX2, VEGF) are characteristic to HGSOCs. We discerned dysregulations (expressions/activity) in SREBP2, SRB-1, STAR, and HMGCR along with Akt/pAkt^Thr308^, mTOR/pmTOR^Ser2448^, p70S6K, P38MAPK, HIF-1α, COX2, and VEGF proteins within high cohort. Herein, poorly differentiated tumors with escalated HMGCR activity overproduced cholesterol thereby rigidifying their cell membranes to restrain Pt-DNA adduct retention. With a carboplatin IC_50_ of 5.23µM, high cohort tumors generated lesser drug-induced ROS and espoused unaltered mitochondrial-membrane depolarization and DNA damage profiles. These parameters were moderately altered in the borderline-HGSOC cohort possessing relatively less rigid membranes and a lower carboplatin IC_50_ of 2.78µM. Accordingly, borderline and high cohorts were respectively denoted as intermediate responder and non-responder of carboplatin. On the contrary, the cholesterol-deficient optimum cohort (IC_50_-1.59µM) with fluid membranes was a carboplatin responder group.

Our study established the candidature of abnormal cholesterol and PI3K/Akt/mTOR (protein-level) statuses as predictive markers to screen HGSOCs for carboplatin responses before therapy.

## INTRODUCTION

Increasing High-Grade Serous Ovarian Cancer (HGSOC) incidences, relapse, recurrence, and mortalities have helmed it as a ‘global clinical concern’ [1, 2]. Late presentations and non-specific screening with Cancer antigen-125 (Ca-125) estimation have been the pivotal contributors to this scenario [3–7]. Despite the conventional practice of ‘first line of treatment’ with platinum-ligated drugs like ‘carboplatin’ alongside cytoreductive/primary debulking surgeries, HGSOC patients shortly relapse within six months of therapy completion [8–11]. Hence, rationalizing chemotherapy regimens among patients is more important than following the convention without prior knowledge of its outcomes.

Recently, the characterization of heterogeneous HGSOC tumor microenvironment (TME) is relied on for identifying ‘predictive biomarkers’ that potentiate the categorization of HGSOC subjects as ‘carboplatin responders’ and ‘non-responders’ before therapy [12, 13]. In this regard, HGSOC-related ‘cholesterol metabolism reprogramming’ along with the underlying ‘PI3K/Akt/mTOR dysregulation’ can provide clues for interpreting carboplatin treatment fates [14–20]. During malignant transformation, steroidogenic ovaries fuel uncontrolled cellular proliferation by altering their cholesterol metabolism and the underlying signaling milieu. Serine-threonine kinases like Phosphoinositide3 Kinase (PI3K), Protein Kinase B (Akt), and mammalian Target of Rapamycin (mTOR) are the pivotal regulators of these signaling cascades directly responsible for metabolic reprogramming of HGSOC tumors [21–27]. Therefore, comprehending the chemico-biological attributes of HGSOC subjects can elucidate their relative sensitivities and responsiveness towards therapeutic agents.

Hyperactivated PI3K/Akt/mTOR pathway reportedly drives anomalous cross-talks of the prosurvival effectors namely P70 Ribosomal protein S6 kinase (P70S6K1), P38MAPKinase (P38MAPK), Hypoxia Inducible Factor-1α (HIF-1α), Cyclooxygenase 2 (COX2), and Vascular Endothelial Growth Factor (VEGF) in its downstream [28–31]. Such a prosurvival molecular interplay furthermore orchestrates endogenous and exogenous effectors deciding cholesterol turnovers within the TME. mTOR activation, for instance, drives the nuclear translocation of Sterol Response Element Binding Protein 2 (SREBP2) to promote the expression of 3-Hydroxy-3-methylglutaryl-coenzyme A Reductase (HMGCR), the rate-limiting enzyme of cholesterol synthesizing ‘mevalonate pathway’ [32–49]. Heightened PI3K/Akt/mTOR signaling also evokes the overexpression of Scavenger Receptor Class B Type 1 (SRB1) and Steroidogenic Acute Regulatory (STAR) proteins in HGSOC tumors [33–38]. Through SRB1, HGSOC cells exogenously re-route cholesterol from blood and tissue fluids for metabolizing it in the mitochondria following export by STAR [45, 48]. Hence, HGSOC cells maintain their cellular repertoire of cholesterol by employing endogenous synthesis or exogenous import to suffice their metabolic needs during malignant transformation [49]. In either case, PI3K/Akt/mTOR activity synergizes cholesterol metabolism reprogramming to bestow HGSOCs with an aggressive phenotype that paves the way for intrinsic or acquired carboplatin resistance [39, 40]. For example, PI3K/Akt/mTOR mediated enhancement of cholesterol turnover rigidifies cell membranes to hinder drug import for reducing intracellular accumulation of chemotherapeutics like carboplatin. Again, hypoxia eventuates with cholesterol metabolism acceleration in HGSOC TME, stimulating COX2, HIF-1α, and VEGF expressions which are also a characteristic of carboplatin-resistant tumors [32–33]. Such an intertwining abnormalcy in cholesterol metabolism and PI3K/Akt/mTOR signaling envisages crucial molecular anecdotes decisive for carboplatin treatment efficacy among HGSOCs.

Based on these facts, this present ‘pilot study’ was proposed to explore the credibility of the ‘protein expression’ statuses of HGSOC-related PI3K/Akt/mTOR candidates and cholesterol biosynthesis regulators along with distinct ‘blood-cholesterol profiles’ in predicting carboplatin therapy response.

## MATERIAL AND METHODS

### Patient Study

Following ethical clearance (IEC Ref: CNCI-IEC-SM-2022-34), a sample of 31 Non-NACT HGSOC cases was recruited for this study in accordance with Flat Pilot Study Sample Size Rules of Thumb and compared with 12 other gynaecological cancer cases and 18 normal volunteers[50]. Over six months, clinically confirmed HGSOC cases were recruited as study subjects from the ovarian cancer patients visiting the outpatient clinic (OPD) of Chittaranjan National Cancer Institute (CNCI), Kolkata, INDIA. Non-NACT HGSOC subjects (age: 30-65yrs; ECOG status: 0-1) with no history of anthracycline, vinorelbine, aspirin, statin, or radiation treatment and concurrent dyslipidemia in addition to ‘primary debulking’ surgery recommendation were included while smokers, alcohol intakers, and pregnant/nursing women were excluded. This pilot study was observational with no extension till follow-up.

### Sample Collection and Processing

Following the acquisition of signed Informed Consent Forms, tissue samples from individual patients were collected post-surgery while their blood samples were obtained a day before the operation. Patients with other gynaecological cancers (cervical, endometrial, vulvar) with hysterectomy recommendations and normal non-cancer volunteers were also included as references. Blood samples were collected from all three groups for cholesterol quantification which formed a basis for the comparative categorization of HGSOC patients into ‘optimum’, ‘borderline’, and ‘high’ cholesterol cohorts. Tissue cholesterol levels of HGSOC samples were estimated in comparison with normal ovaries collected from other gynaecological patients cohort-specifically. The blood cholesterol range referent for platinum therapy outcomes in HGSOC patients was thereafter decided by their relative tissue cholesterol titers, molecular deregulations, and carboplatin response. Sample collection and processing were undertaken as per the norms of the Institutional Human Ethics Committee, CNCI, Kolkata, INDIA.

### Cell culture

SKOV3, an ovarian epithelial cancer cell line, and its resistant counterpart SKOV3^R^ were respectively maintained in proportionate volumes of RPMI and DMEM supplemented with 10% FBS and antibiotics (gentamycin 40 μg, penicillin 100 units, and streptomycin 10 μg/ml) at 37°C in a humidified CO_2_(∼5%) incubator. Normal ovarian cell line IOSE was maintained in equal volumes of MCDB-105 and Medium-199 (v/v) with the same supplements in similar conditions.

### Ex vivo Primary Culture

Surgical tissue (3-optimum, 5-borderline, 10-high) samples transferred in chilled Phosphate Buffered Saline (PBS; pH-7.4) immediately after collection were washed well in sterile conditions within Laminar Airflow Hood (Klenzaids). Tissue samples were minced in chilled PBS using sterile fine forceps and scalpel followed by their transfer in microcentrifuge tubes containing Type-I Collagenase (HIMEDIA) prepared in DMEM-F12 medium supplemented mildly with ~4% FBS. Tissue fragments were subjected to a controlled slow digestion process for 4-6 hrs at 37°C in a humidified CO_2_ (∼5%) incubator with interim shaking for dislodging digested cells. The single cells were collected by centrifuging these tubes at 3000rpm for 6 min at room temperature (RT). After discarding the supernatant, the resulting pellets were washed once in PBS by centrifugation at 1500 rpm for 5 min followed by resuspension in DMEM-F-12 supplemented with 20% FBS. All cell suspensions were plated in 55mm culture plates and observed for cell growth. Primary plating had gentamycin (5μg/ml) and streptomycin (5μg/ml) added to the culture medium to prevent initial chances of contamination. Exponential cell growth was achieved by 3^rd^ day of plating upon transfer to primary plating-derived conditioned medium along with fresh DMEM-F-12 medium (1:4) containing 15% FBS. This condition was maintained with a gradual reduction of FBS proportions to 10% until large viable colonies grew after the 5^th^ day. At about the 7^th^ day, the first passage was given with mild trypsinization followed by bulk culturing of cells for about 14 days. *Ex vivo* cultures were finally maintained in antibiotic-free DMEM-F12 medium supplemented with 10% FBS up to 3-4 passages. Cells were harvested for carboplatin-response assessment intermittently.

### Development of Carboplatin-resistant Cell line

SKOV3 cells were subjected to treatment with a broad logarithmic range of carboplatin doses ranging from 0.1 μM to 10μM. By MTT assay, the IC_20_, IC_30_, IC_50_, and IC_70_ carboplatin doses were calculated. As per Mahapatra et al., 2022, SKOV3 cells were subjected to ‘pulse treatment ‘for the development of a carboplatin-resistant subline SKOV3^R^ within 3 months [20].

### MTT Assay

Ahead of initiating ‘pulse treatment’, admissible carboplatin doses were selected by MTT assay following Mahapatra et al., 2022 [20]. Establishment of SKOV3^R^ was compared for its carboplatin response against its parental counter-part SKOV3 by calculating fold resistance [Fold Resistance= (IC_50_ of SKOV3^R^)/ (IC_50_ of SKOV3^R^)] value after MTT Assay. A similar protocol was also followed for separately identifying the carboplatin dose response for *ex vivo* HGSOC cell cultures from optimum, borderline, and high cohorts after treatment with a logarithmic concentration range (0.1-100μM) of carboplatin. Results were enumerated as Semi-log graphs and finally represented in terms of cellular survival (%) trend curve.

### Blood Plasma Isolation

Equal volumes of Histopaque (SIGMA) and blood were carefully taken within 15ml tubes and centrifuged at 400g for about 30mins. After centrifugation, this mix was separated into three distinct phases of plasma, peripheral blood mononuclear cells (PBMC), histopaque, and lysed-RBCs. Plasma was collected in microcentrifuge tubes from the uppermost clear phase and stored at −20°C until use for cholesterol quantification.

### Tissue and cell lysate preparation for Cholesterol Quantification

Fresh unfixed tissue samples were homogenized well using Dounce Homogenizer in Cholesterol Assay Buffer provided with the Cholesterol Quantification Kit (abcam). Additionally, harvested single cells from primary cultures and cell lines (SKOV3, SKOV3^R^, IOSE) were pelleted down in microcentrifuge tubes following centrifugation at 1500 rpm for 5 mins. These pellets were further resuspended in Cholesterol Assay Buffer for lysis. The resulting lysates were cold-centrifuged at 13000g for 10mins followed by the collection of supernatants in fresh tubes which were stored at −20°C until use for cholesterol quantification.

### Cholesterol Quantification

Quantitative estimation of cholesterol in the plasma and tissue lysates was undertaken with the aid of a commercially available Cholesterol Quantification Kit (abcam) as per the provided protocol. Accordingly, 50μl of samples respectively constituted by 1μl of plasma or tissue/cell lysates along with 49 μl of Cholesterol Assay Buffer were added to 96well plates pre-filled with 50μl Cholesterol Reaction Mix (44 μl Cholesterol Assay Buffer, 2 μl Cholesterol Probe, 2μl Enzyme Mix, 2 μl Cholesterol Esterase Enzyme). Total reaction mixtures (100μl) were incubated at 37°C for 1h followed by spectrophotometric recording of absorbances at 570nm. Cholesterol levels were calculated after interpolating the recorded absorbances in a standard curve followed by representation in mg/dl. All experiments were repeated thrice in light protected conditions and values were expressed as Mean ±S.D.

### Histopathological Study

Freshly obtained HGSOC samples were washed in ice cold PBS (pH-7.4) and fixed in Neutral Buffered Formalin (NBF) for 48h and processed for histopathological analysis after the standardized protocol of Mahapatra et. al, 2022 [20]. Haematoxylin and Eosin stained slides were mounted with DPX for microscopic analysis. Each slide was scanned for at least 20 fields to identify and locate histopathological changes under a light microscope (Zeiss).

### Retrospective Analysis

In order to select relevant molecular markers for study, The Cancer Genome Atlas (TCGA) database was referred. TCGA reported 28-HGSOC cases were thoroughly mined for the enlisted genetic markers which presented high mutation or copy number variation frequencies. Accordingly, PIK3CA, AKT1, mTOR, RPS6KB1, MAP3K1, HIF1A, PTGS2, VEGFA, SREBPF1, SREBPF2 and STAR were identified. Proteins encoded by these genes-PI3K, Akt, mTOR, p70S6K, P38MAPK, HIF-1α, COX2, VEGF, SREBP1, SREBP2 and STAR were selected as study candidates in addition to the protein SRB-1. Despite no reports of SCARB-1 which encodes for SRB-1, a deliberate inclusion of SRB-1 was made because of its function that suffices the study objectives. Recorded observations were represented in the form of a Heat Map prepared with OncoGrid.

With the help of KM Plotter, the dependency of patient survival upon SREBP1, SREBP2, SRB-1, and STAR expressions was studied. The online platform, Gene Expression Profiling and Interactive Analysis (GEPIA) was referred for analyzing the significance of conjugative expressions of these cholesterol metabolism regulators in ovarian cancer scenarios. Additionally, DepMap Portal was investigated for the survival dependency of ovarian cancer cell line SKOV3 upon SREBP1, SREBP2, SRB-1, and STAR.

### STRING Database Analysis

Interactions of the selected proteins were verified further in STRING Database. Results were represented in the form of available interactome-diagrams depicting protein-clusters with numerical designations. These included Akt cluster[Akt1(1); Akt2(2); Akt3(3)], P38MAPK cluster [MAPK11(4); MAPK14(5); MAPK13(6); MAPK12(7)] and mTOR cluster [Akt1S1(8); EIF-4EBP1(9); RPTOR(10); MLST8(11); MTOR(12)] interacting with P70S6K [RPS6KB1(13)], COX2[PTGS2(14)], HIF1α[HIF-1A(15)], VEGF [VEGFA(16)], STAR[STAR(17)], SREBP2[SREBF2(18)], SREBP1[SREBF1(19)] and SRB-1[SCARB(20)]. All interactions were mapped based on significant interactions only.

### Tissue and Cell Immunofluorescence

Paraffin-embedded HGSOC tissue sections (∼5μm thick) were analyzed for the locations of SREBP2, SRB-1, and STAR in relation to SREBP1 by multiplexing. For this purpose, paraffin from stretched tissue sections was removed by heating at 65°C for 20mins followed by xylene treatment and rehydration through 100%, 90%, 70%, and 50% alcohol downgrades. After serial washing (5 times) of these sections in PBS for 10min, ‘antigen-retrieval’ was carried out using pre-heated Citrate Buffer [pH-6] at 85°C for 20mins followed by three PBS washes. These sections were incubated overnight with respective primary antibodies diluted in 1% BSA solution (1:1000) within a humidified chamber at 4°C. Excess primary antibodies were washed in 1X PBS on the following day and the slides were further incubated with fluorophore-conjugated secondary antibodies (1:500) in 1% BSA solution for 2h at room temperature followed by counterstaining with 4’,6-diamidino-2-phenylindole (DAPI), a nuclear stain. These slides were dehydrated through successive alcohol upgrades and xylene. Finally, they were mounted in DPX for observation under a laser confocal microscope (Zeiss).

SKOV3 and SKOV3^R^ cells were seeded (2.5 × 10^5^cells/well) onto coverslips placed within 6-well plates which were fixed and immunostained as per Mahapatra et. al, 2022 [20]. All fluorescent micrographs of tissue sections and cells were represented as composite images along with those recorded separately for DAPI, APC, and FITC filters.

### Isolation and Estimation of Proteins

HGSOC samples were washed, processed, and pooled separately. Tissue pieces were dried, weighed, and homogenized in Radio-Immunoprecipitation Assay Lysis buffer (RIPA; pH-8 comprising of 5M NaCl, 0.5M EDTA,1M Tris, NP-40,10% Sodium Deoxycholate,10% SDS). All extracts were ice incubated for 30 min followed by sonication and centrifugation at 10,000g for 20 min at 4°C. The resulting supernatants were spectrophotometrically (VARIAN) evaluated for their protein contents using 1X Bradford’s reagent against a BSA standard curve. Absorbances were recorded at 595 nm with the experiment being repeated 5 times. Protein concentrations were represented in µg/µl.

### Western Blot Analysis

Protein expressions of selected study candidates were comparatively ascertained by western blotting. Equitable amounts of tissue protein (~30μg) were respectively loaded into each well of SDS-polyacrylamide gels, electrophoretically separated following standardized laboratory protocol [20]. Chromogenic substrate 5-bromo, 4-chloro, 3-indoylphosphate/ Nitro-Blue tetrazolium (BCIP/NBT) was used for visualizing protein expressions in the form of bands. β-actin was used as a loading control protein. Respective band intensities were calculated in ImageJ after normalization against loading control. All experiments were performed in triplicate.

### Co-Immunoprecipitation

Interactions of SREBP2 and its probable associates were ascertained by co-immunoprecipitation assay. To the microcentrifuge tubes containing 200μg of protein samples, protease inhibitors along with SREBP2 antibodies (2μg/100-200μg of protein) were added. All tubes were thereafter loaded into a tube rotator which was kept in a 4°C cold room overnight to enable immunoprecipitation of SREBP2-associated protein complex. Next up, the tube contents were transferred to columns containing filters coated with Protein-A Sepharose beads which were further kept at 4°C overnight under constant shaking. On the following day, non-specific proteins were washed off the cell lysates in 1X and 0.1X IP Buffer respectively by cold-centrifugation at 12000g for 1min. The purified protein complex was thereafter eluted out using IP Elution Buffer followed by cold-centrifugation at 12000g for 1min. IP elutes were qualitatively analyzed for expression of interacting protein candidates by western blotting.

### HMGCoA Reductase Activity

Quantitative estimation of HMGCoA Reductase activities in tissue and cellular proteins was performed with a commercial kit (abcam) as per the manufacturer’s instructions. Lovastatin, the provided HMGCR inhibitor was dissolved in DMSO and added to the samples along with other co-factors. Reaction mixtures were incubated at 37°C for 10mins in humidified conditions. Absorbances were recorded spectrophotometrically at 340nm in three kinetic cycles. Results were enumerated in terms of units of enzyme (*10) inhibited after interpolation in a stand-curve using GraphPad Prism software. Experiments were repeated thrice.

### In silico Analysis

The bioinformatics clustering algorithm ClusPro was used for protein-protein docking. Three-dimensional structure of SREBP2 (PDB-ID: 7AZN) in conjugation with mTOR (PDB-ID: 7PEC), p70S6K (PDB-ID: 3A60), HIF-1α (PDB-ID: 1H2M), SRB-1(PDB-ID: 5KTF) and STAR (PDB-ID: 2R55) were computed for their interaction qualities. With a clustering-based approach, the conformational spaces’ of interacting proteins were explored for all the possible ‘docking conformations’ or ‘binding-modes’ generated on the basis of structural similarities. In consideration of electrostatic interactions, van der Waals forces, and desolvation effects, these docking conformations were scored for their binding energies to determine the most stable interacting conformations. Energy scores for each cluster were provided by Piper, a Fast Fourier Transform (FFT) based rigid docking program. Clusters with the lowest binding energies depicted the most stable interactions.

### Antigen Staining by flow cytometry

Patient-derived primary cells harvested in PBS supplemented with 10% FBS from 55mm plates after trypsinization were pelleted down by centrifugation at 1500rpm for 6min. Chilled acetone (200μl) was added to these pellets and incubated at −20°C for fixation and permeabilization. Acetone removal was thereafter undertaken by washing and centrifugation of the fixed cells in PBS at 1500rpm for 6min. Fixed cells were blocked in 10%FBS at 4°C for 45min following which they were centrifuged and washed of excess FBS as well at 1500rpm for 6min. Cell pellets were resuspended in PBS supplemented with respective primary antibodies (1:1000) and kept under constant shaking at 4°C overnight. On the following day, a proportion of cell tubes were taken up for antigen staining where cell pellets were further incubated with fluorophore-tagged secondary antibodies at room temperature after the removal of excess primary antibodies for 2h at room temperature. Cells were washed of excess antibodies and thereafter taken up for FACS analysis. Percentage of positively stained cells was represented in Heat Maps generated using the Morpheus Online Application. Mean Fluorescence Intensities (MFIs) were individually represented with the aid of Radar Charts. For double stains, MFIs were represented as the ratio of MFI of pAkt^Thr308^ or SREBP1 versus other antigens. Experiments were repeated twice.

### JC-1 staining

Primary cells were comparatively assessed for their mitochondrial transmembrane potential by staining with JC-1 mitochondrial stain. As per requisite, these cells were incubated with JC-1 staining solution (100μl stain /ml of culture medium) at 37°C for 30 min followed by centrifugation at 400g. Cell pellets were washed twice in 2ml of Assay Buffer. Following aspiration of the supernatant, the excess stain was removed by final washing in 1ml Assay Buffer. These cells were thereafter taken for FACS analysis. All experiments were repeated thrice with results being represented as scatter plots depicting the shift of cellular populations from red and green double-positive quadrant to green positive quadrant. Additionally, results were represented as Red to Green ratio.

SKOV3 and SKOV3^R^ cells were similarly harvested and stained with JC-1 for FACS analysis. All results were represented graphically as the ratio of red to green cell percentages.

### Flow Cytometric Quantification of ROS

Quantitative estimation of ROS was performed by flow cytometry (BD LSR Fortesa; BD Biosciences) in primary cultured *ex vivo* cells as well as SKOV3 and SKOV3^R^ using 2’,7’-dichlorofluorescein dihydroacetate (DCFH-DA; Santa Cruz) as per laboratory protocol [20]. Scatter plots and histograms were generated in replicates using Cell Quest software. Results are represented as overlay histogram curves generated by Flowjo software.

### Quantification of intracellular Pt-DNA adduct retention by flow cytometry

Carboplatin treated primary cultured cells were harvested and pelleted down by centrifugation at 1500rpm for 5min followed by acetone fixation at −20°C for 20min. After acetone removal by centrifugation, fixed cells were incubated with an antibody specific to Pt-modified DNA overnight at 4°C. Thereafter, excess antibodies were washed away by centrifugation at 1500rpm for 5min followed by incubation with FITC conjugated secondary antibodies for 2h at room temperature followed by flow cytometric detection. Results were represented as histograms. Experiments were repeated thrice.

### Comet Assay

The clastogenic effect of carboplatin on DNA was assessed by comet assay (single cell gel electrophoresis). Concisely, a suspension of 0.6% (w/v) low melting agarose (LMA; Sigma-Aldrich, USA) and isolated primary cells was smeared over a frosted microscopic glass slide which was priorly coated with 0.75% (w/v) normal melting agarose (NMA; Lonza, USA). Following solidification at 4°C, cell and nuclear membranes were lysed in lysis buffer (pH 10). Exposed DNA from the lysed-out leukocytes was unwound in a highly alkaline electrophoresis buffer prior to electrophoresis for 20 min (300 mA, 20 V). Slides were washed in neutralizing buffer thrice, stained with ethidium bromide (final concentration 40μg/ml), and examined under a fluorescence microscope (Leica). Image analysis, head DNA quantification, comet tail DNA length estimation and comet tail moment calculation were performed using Komet Software.

### Laurdan Staining

Primary Cells, SKOV3, and SKOV3^R^ cells were stained with ‘LAURDAN’ (Invitrogen), a fluorescent stain, and analyzed for their 450/530 ratio in their fluorescent emissions with the help of a spectrofluorometer (VARIAN). On being excited by 350nm laser, the dye fluoresced differently depending upon the rigidities of the cell membrane. With fluid membranes, fluorescence emissions were detected at around 500nm while with rigid membranes this was found to be around 450nm. All intensities were represented graphically with experiments being repeated thrice. Results were expressed as Mean±SD.

### Statistical Analysis

Mean values of the data were compared by factorial Analysis of Variance (ANOVA). The relationship between the studied parameters was analyzed by calculating Pearson’s correlation coefficient using Prism GraphPad Software. Band Intensities of western blot were calculated using Image J software. All data were expressed as mean ± standard deviation (S.D.). p value calculations were performed using Prism GraphPad Software where ***p < 0.0001 and **p<0.05 were considered statistically significant.

## RESULTS

### High-Grade Serous Ovarian Carcinoma envisages Aberrant Blood and Tissue Cholesterol Profiles

The present study deciphers the clinical relevance of reprogrammed cholesterol metabolism in predicting the chemotherapy response of HGSOC towards carboplatin. We initiated the study by identifying if aberrant cholesterol metabolism was an issue for the HGSOC scenario (Fig. 1a and S1a). Over six months, three human study groups- a) HGSOC (n=31), b) Other gynaecological cancer subjects (vaginal tumor/endometrioid cancer/ vulvar tumor; n=12), and c) Normal Non-Cancer Volunteers (n=18) were recruited randomly. With the later groups as references, a comparative plasma cholesterol profile for Non-NACT HGSOC cases with recommendations for primary debulking surgery was quantitatively prepared (Fig. S1a). Accordingly, these respective groups were categorized as ‘optimum’ (<129mg/dl), ‘borderline’ (130-159mg/dl), and ‘high’ (>160mg/dl) cholesterol cohorts (Fig. 1a, S1a, 1b, and S1b). The mean plasma cholesterol levels of the optimal HGSOC cohort was 88.60±8.74 mg/dl which was comparable to that of the reference groups (normal volunteers: 95.870±19.21 mg/dl; other cancers:101.52±16.84mg/dl). In the borderline cohort, HGSOC patients recorded 1.7fold higher (142.965±9.03mg/dl) blood cholesterol levels than the normal volunteers (137.513±12.65mg/dl). This was comparably close to the borderline-other cancer cases (145.265±17.34mg/dl). HGSOC patients from the high cohort presented a distinctly escalated (290.625±13.650mg/dl) plasma cholesterol titer than the normal volunteers (227.115±13.650mg/dl/1.3fold;p<0.05) and other cancer patients (210.672±16.465mg/dl/1.4fold; p<0.0001) as well (Fig. 1b and S1b). The distribution of these three study groups among the optimal, borderline, and high blood cholesterol cohorts was further analyzed. As depicted in Fig. 1c, 54.84% of HGSOC patients, 50% of other cancer patients, and 33.33% of normal volunteers could be clustered in the high cohort. The borderline cohort was comprised of about 32.25% HGSOC cases alongside an equitable 16.67% of other cancer and normal volunteer candidates. Major contributors of the optimal cohort were identified as either normal volunteers (~50%) or other cancer subjects (~16.67%) followed by the HGSOC patients (~12.9%) in the minority. Therefore, a clear predominance of subjects with high or borderline blood cholesterol levels was noticed among the HGSOC study group (Fig. 1c). These findings associated cholesterol aberrance with the HGSOC scenario.

**Fig. 1:**
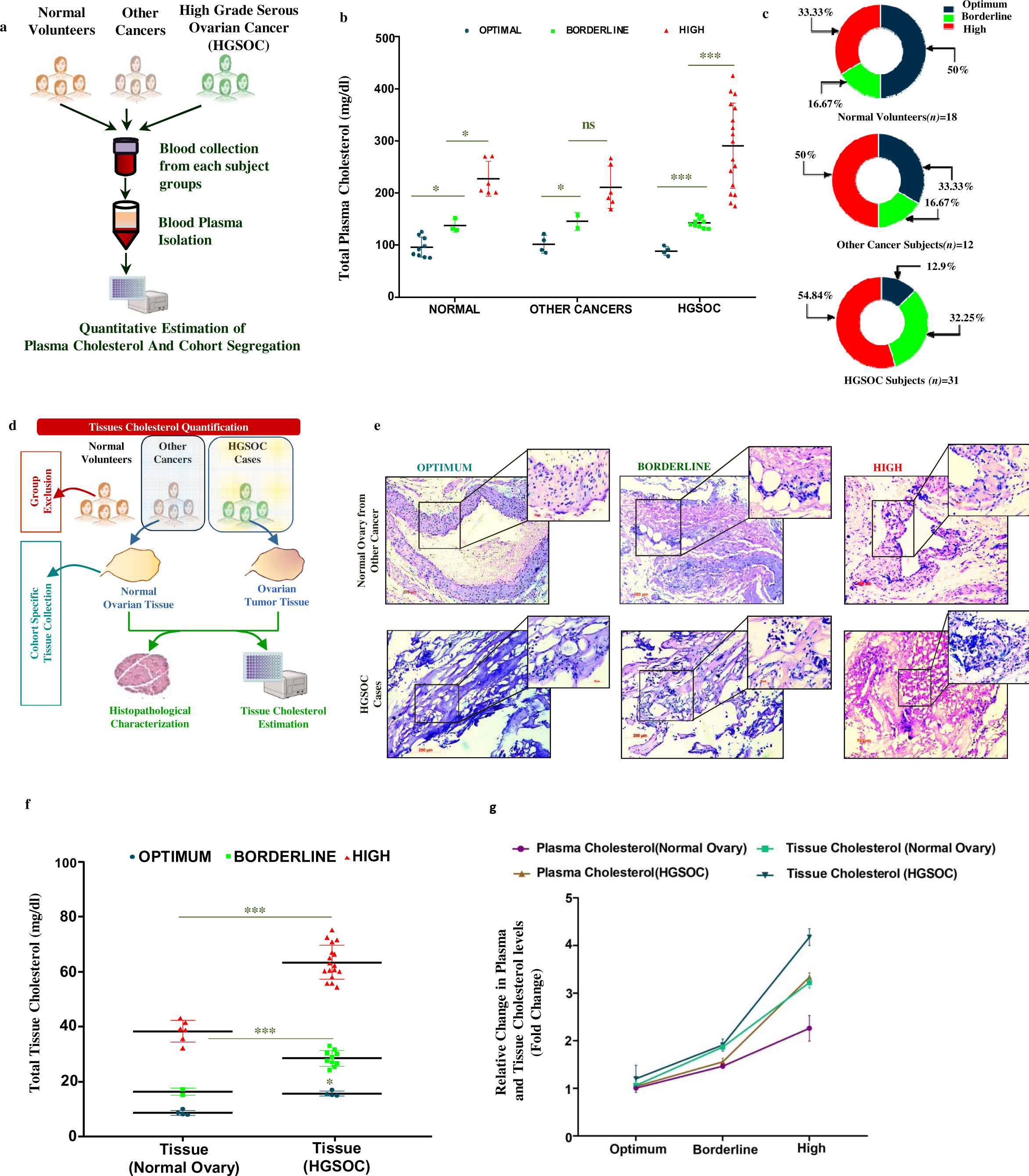
Aberrant Blood and Tissue Cholesterol Profiles are intrinsic to HGSOC: (a) An Overview of Work-Plan followed while quantifying cholesterol levels in the blood plasma of HGSOC patients, other cancer subjects and normal volunteers. (b) A graphical anecdote of individual blood cholesterol values categorizing the three human-study groups into optimum, borderline and high cohorts respectively. (c) Pie-charts representing cohort wise distribution patterns of normal volunteers, other cancer and HGSOC subjects. (d) Study design for evaluating tissue cholesterol levels in HGSOC and other cancer patients. (e) Micrographs delineating cohort-specific remodelled histology of HGSOC tissues and normal ovaries [Main image-100X (Scale bar: 200µm); Insets-400X (Scale bar: 50µm)]. (f) Scatter plots representing cohort-specific tissue cholesterol profiles of HGSOC and normal ovarian tissues. (g) Relative trends in blood and tissue cholesterol alterations among HGSOC and other cancer cohorts. Error bars in (b), and (f) represents standard deviation of mean values of each replicates with *p<0.05; ***p<0.0001 being considered significant.

Next up, the influence of such an abnormal blood cholesterol profile upon its relative abundance within malignant ovarian tissues was assessed. For this purpose, the cholesterol content of histopathologically annotated normal as well as HGSOC ovarian samples obtained respectively from optimal, borderline, and high blood cholesterol cohorts was quantified (Fig. 1d and S1a). Hysterectomically resected ovarian tissues from other cancer subjects were included as normal reference after histopathological confirmation. Henceforth, the normal volunteers group was rationally omitted (Fig. 1d and S1a). Results in Fig. 1e, 1f, and S1c indicated that HGSOC tissues of the optimal cohort possessed vacuolation with papillary extensions and contained a cholesterol titer of 15.65±0.675mg/dl. Conversely, the normal ovarian samples with a cholesterol content of 8.86±0.612mg/dl were non-dysplastic. Herein, the HGSOC tissue cholesterol was 1.78fold (p<0.05) greater compared to normal ovaries in the optimal cohort (Fig. S1d). On the other hand, borderline HGSOC tissues with a cholesterol abundance of 28.48±0.88mg/dl appeared to be cribriform with poorly differentiated bizarre structured cells. Correspondingly, the normal ovaries with 16.29±1.01mg/dl of cholesterol quantitate exhibited a solid well differentiable architecture (Fig. 1e, 1f, and S1c). Yet again, the borderline HGSOC tissues emerged to be more cholesterol-enriched (1.74folds; p<0.05) than the normal (Fig. S1d). Poorly differentiated HGSOC tissues abundant in mononuclear infiltrates along with cells having dense eosinophilic cytoplasm and large conspicuous nuclei were noteworthy among high cohort (Fig. 1e and S1c). These tissues were quantified with an overtly escalated cholesterol level (63.418±6.291mg/dl/1.6 fold; p<0.0001) unlike the normal ovaries (38.3±3.983mg/dl) lacking any abnormal histopathological anecdotes (Fig. 1e, 1f, S1c, and S1d). Once again, these observations affirmed the presence of a reprogrammed cholesterol metabolic profile in aggravated malignancies like HGSOC.

To strengthen these observations, a cohort-based comparative cholesterol profile for blood and tissues of other cancer and HGSOC groups was designed (Fig. 1g). Cholesterol titers in the plasma of high HGSOC cases were calculated to be 1.4 fold and 1.6 fold higher compared to the borderline and optimal HGSOC subgroups. Likewise, HGSOC tissue cholesterol profiles were also escalated by 1.8 and 4 folds respectively. A collateral increase in the blood and tissue cholesterol levels was quite evident among HGSOC cases (Fig. 1g). Contrarily, normal ovaries from the high cohort recorded 1.75 fold and 2.35 fold higher cholesterol levels than those from the borderline and optimal cohorts (Fig. 1g). Increasing trends in blood specific cholesterol titers did not align with that of the normal tissues. Hence, a prominent rise in the plasma, as well as tissue cholesterol levels, was typically appropriated for HGSOC cases. These findings conclusively established that cholesterol aberrance was an exclusive occurrence for the HGSOC scenario in this pilot study.

### HGSOC Tissues with Heightened Cholesterol Levels display Dysregulation in the Protein Expression Status of PI3K/Akt/mTOR Candidates and Cholesterol Metabolism Regulators

As aforementioned, dysregulation of prosurvival PI3K/Akt/mTOR signaling expedites cholesterol biosynthesis in HGSOC TME to direct therapy evasion. Hence, characterization of HGSOC TME for cohort-specific molecular signatures was preferred ahead of assessing carboplatin therapy response (Fig. 2a). In order to identify credible molecular markers for study, the TCGA database was explored thoroughly. Herein, 28 HGSOC cases with several deregulated molecular markers were reported. As shown in the Heat map of Fig. 2b, only the prime PI3K/Akt/mTOR signaling effectors intricately regulating intracellular cholesterol biosynthesis, uptake, and metabolism were selected as study candidates based upon high mutation or copy number variation (CNV) frequencies. These included 8 ‘prosurvival effectors’ (PIK3CA/PI3K, AKT1/Akt, mTOR/mTOR, RPS6KB1/RS6K, MAP3K1/p38MAPK, HIF1A/HIF-1α, PTGS2/COX2, VEGFA/VEGF) and 3 ‘cholesterol metabolism regulators’ (SREBPF1/SREBP, SREBPF2/SREBP2, STAR/STAR) (Fig. 2b and S2a). Regardless of insignificant reports (Fig. S2a), the inclusion of SRB-1 protein (*SCARB-1*gene) was deliberated because of its role in facilitating nonendocytic cholesterol uptake from exogenous repertoires (blood/tissue fluids). In a function dependent manner, these candidates were further categorized as ‘upstream prosurvival effectors’, ‘downstream prosurvival effectors’, and ‘cholesterol metabolism regulators’ for experimental and data analysis conveniences (Fig. 2c). Thereafter, STRING Database was referred for deciphering the prevalence of any possible protein level interactions between these selected candidates. Interactome map in Fig. 2d delineated that the prosurvival effectors and cholesterol metabolism regulators interacted closely with high confidence interval values which thereby approved the marker selection strategy (Fig. 2d). Thus, the signaling milieu responsible for the cohort-specific phenotypic heterogeneity of HGSOC TMEs was comprehended through protein expression and interaction studies.

**Fig. 2:**
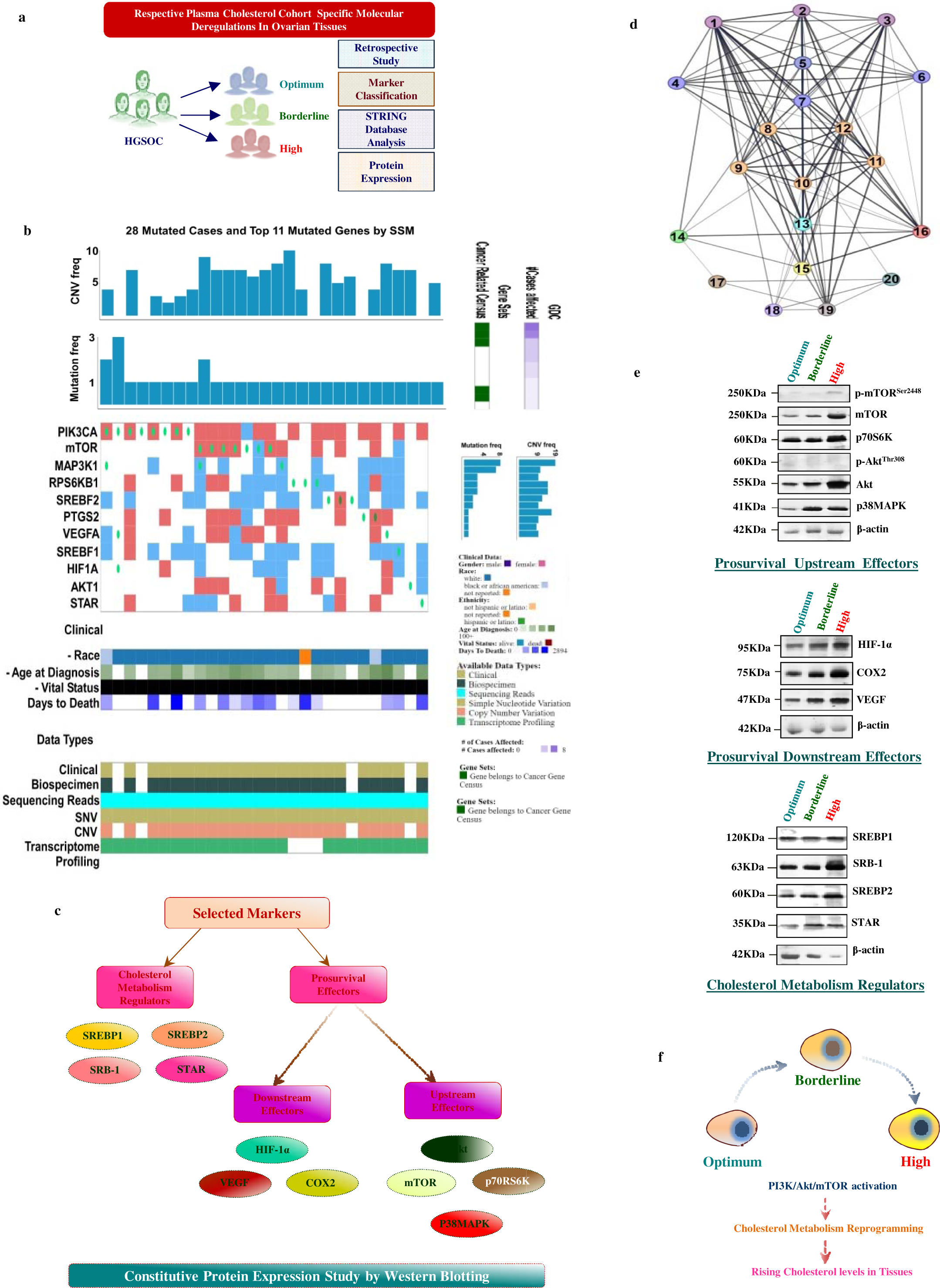
HGSOC tissues with enhanced Cholesterol turnover displayed upregulated PI3K/Akt/mTOR candidates : (a) Schematic representation of the study design. (b) Heat map portraying comparative statuses of TCGA cases reporting markers with high mutation and copy number variation frequencies. (c) Classification of selected study candidates into upstream and downstream prosurvival effectors in addition to cholesterol metabolism regulators. (d) STRING Database Interactions involving Akt cluster[Akt1(1); Akt2(2); Akt3(3)], P38MAPK cluster [MAPK11(4); MAPK14(5); MAPK13(6); MAPK12(7)], mTOR cluster [Akt1S1(8); EIF-4EBP1(9); RPTOR(10); MLST8(11); MTOR(12)], P70S6K [RPS6KB1(13)], COX2[PTGS2(14)], HIF1α[HIF-1A(15)], VEGF [VEGFA(16)], STAR[STAR(17)], SREBP2[SREBF2(18)], SREBP1[SREBF1(19)] and SRB-1[SCARB(20)]. (e) Western blots showing cohort-specific expressions of cholesterol metabolism regulators, downstream and upstream prosurvival effectors. (f) Schematic portrayal of HGSOC TME remodelling by PI3K/Akt/mTOR mediated activation of cholesterol metabolism regulators.

Serial assessment of ‘upstream prosurvival effectors’, ‘downstream prosurvival effectors’, and ‘cholesterol metabolism regulators’ at protein levels was undertaken by western blot analysis (Fig. 2e). Herein, significantly elevated expressions of specific upstream prosurvival effectors-mTOR (5.38 fold; p<0.0001), RS6K (5.33 fold; p<0.0001), Akt (5.76 fold; p<0.0001), and P38MAPK (2.93 fold; p<0.0001) was noteworthy among high cohort HGSOC tissues compared to the borderline. Again, the borderline subjects presented hiked expression profiles of the same in relation to the optimum cohort (Fig. 2e and S2b). Activated forms of Akt and mTOR i.e., pAkt^Thr308^ and pmTOR^Ser2448^ emerged with poor expressions and relative band intensities across all cohorts (Fig. 2e and S2b). Correlatively, the expression of downstream prosurvival effectors was examined to ascertain if this PI3K/Akt/mTOR activation in the upstream transcended downwards as well. Unlike the optimal cohort, HIF-1α (borderline: 4.99 fold; high: 5.648 fold; p<0.0001), VEGF (borderline: 3.88 fold, high: 4.52 fold; p<0.0001), and COX2 (borderline: 4.85 fold, high: 5.25 fold; p<0.0001) were distinctly raised (Fig. 2e and S2c). Hence, expressions of cholesterol metabolism regulators were identified to determine if an activated PI3K/Akt/mTOR could potentiate their upregulation in the borderline and high cohorts especially. As evident from Fig. 2e and S2d, overexpression of SREBP2, SRB-1, and STAR was predominant among high cohort HGSOC patients compared to borderline and optimal cohorts respectively. SREBP2 expression gradually surged by 3.75 (p<0.05) and 4.34 (p<0.0001) folds among borderline and high cohorts compared to optimal. SRB-1 protein was also raised by 3.75 fold (p<0.05) and 5.26 fold (p<0.0001) within borderline and high HGSOC cohorts than the optimal. Unlike SREBP2, SREBP1 expressions remained comparable among all three cohorts (Fig. 2e and S2d). As perfectly portrayed in Fig.2f, PI3K/Akt/mTOR upregulation evokes cholesterol metabolism regulators for overproducing cholesterol among HGSOC tissues.

### Crosstalk between Specific PI3K/Akt/mTOR Pathway Candidates and Cholesterol Metabolism Regulators Enhances Cholesterol Turnover in HGSOC Tissues

The effect of upregulated PI3K/Akt/mTOR candidate expressions upon increased cholesterol turnovers among the HGSOC tissues was further assessed to unravel the mechanism in effect. As per the study design shown in Fig. 3a, patient survival dependency upon the expression profiles of SREBP1, SREBP2, SRB1, and STAR was first investigated using KM Plotter. Poor patient survival with higher expressions of SREBP1 and SREBP2 was apparent among serous grade ovarian cancer subjects treated with platins (Fig. 3b). Survival dependency on SREBP2 was observed to be highly significant (p=0.0001) which was comparable with that of SREBP1 (p=0.0041). Contrastingly, high SREBP2 expressions barely affected the survival of ovarian cancer patients of non-serous categories (Fig.S3b). Patient survivability did not vary with high or low expressions of STAR and SRB-1(Fig.S3b). Furthermore, data collated from GEPIA revealed that SREBP2 and SREBP1 co-expressed among ovarian cancer tissues with a strong positive correlation (r=0.59). On the contrary, SRB-1(r=0.12) as well as STAR(r=0.04) co-expressions with SREBP2 appeared to be poorly correlated among ovarian cancers (Fig.S3a). These results established SREBP1 and SREBP2 as growth essential factors in relation to SRB-1 and STAR in the HGSOC scenario. Therefore, these existent observations were validated by *in silico* analysis using ClusPro’s clustering algorithm platform (Fig. 3c). Herein, all the cholesterol metabolism regulators were subjected to interaction with the upstream and downstream prosurvival effectors *in silico*. As represented in Fig. 3c, only SREBP2 but not SREBP1, interacted with the other two cholesterol metabolism regulators i.e., SRB-1 and STAR along with mTOR, p70S6K, and HIF-1α. With the Piper program, lowest binding energy was calculated for SREBP2 and mTOR (−972.2Kcal/mol) interactions followed by the same with SRB-1(−937.6Kcal/mol), STAR (−856.2Kcal/mol), HIF-1α (−788.9Kcal/mol), and RS6K (−761.6Kcal/mol) (Fig. 3c). Lowest binding energy values indicate spontaneous and strong interactions between the molecules. Accordingly, mTOR, SRB-1, and STAR emerged as the close associates of SREBP2 compared to HIF-1α and RS6K. These findings abided with the String Database observations (Fig.2d) and were suggestive of an inevitable requirement of SREBP2 for HGSOC sustenance. Hence, the signaling interactome was further validated in Co-Immunoprecipitation study where the SREBP2 (IP) complex was immunoprecipitated out of the total tissue protein followed by identification of mTOR, HIF-1α, SRB-1, RS6K, and STAR by immunoblotting (IB). Results depicted in Fig. 3d positively confirmed the prevalence of protein interactions especially within high and borderline cholesterol cohorts unlike the optimal. On this basis, SREBP2’s role as a growth essential factor superseded that of SREBP1 in this study. Additionally, these results were indicative of a probable cholesterol metabolism reprogramming in the HGSOC TME responsible for enhanced tissue cholesterol turnover.

**Fig. 3:**
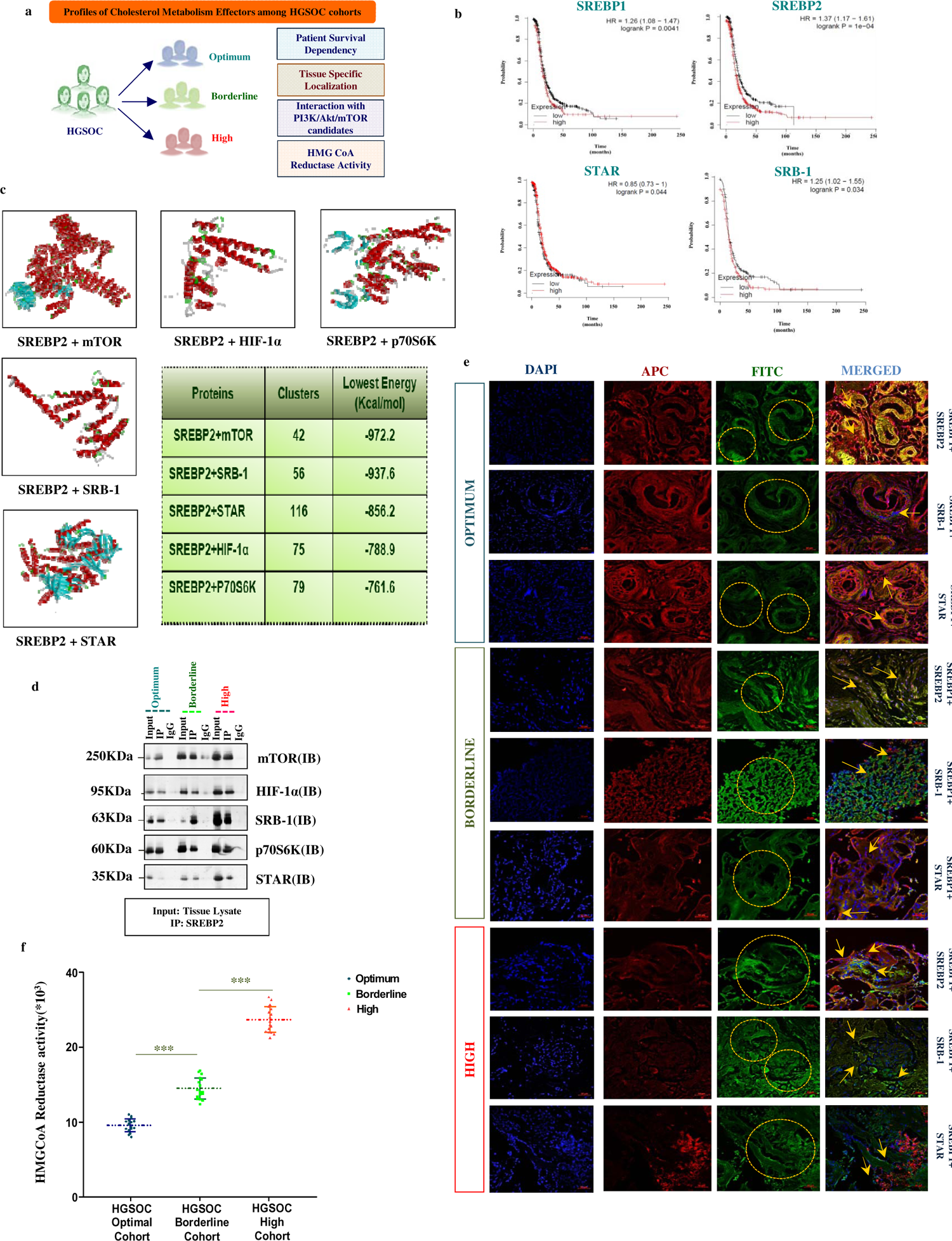
Increased Cholesterol Levels results from exacerbated HMGCR activities promoted by activation of Cholesterol Metabolism Regulators in HGSOC: (a) Diagrammatic representation of the Work plan followed. (b) KM-Plotter data showing survival dependency of serous ovarian cancer patients upon SREBP1, SREBP2, SRB-1, and STAR. (c) *In silico* results delineating plausible interaction between Cholesterol Metabolism Regulators and some key prosurvival candidates of PI3K/Akt/mTOR Pathway. (d) Co-IP studies portraying SREBP2 (IP) interactions with mTOR, HIF-1α, SRB-1, p70S6K and STAR. (e) Tissue IF displaying tissue-specific localization of SREBP2, SRB-1 and STAR with respect to SREBP1 (400X; Scale bar: 50µm). (f) Cohort-wise HMGCR activities being represented as single replicates graphically (Mean± S.D; ***p<0.0001).

Tissue-specific functional disposition of all cholesterol metabolism regulators was further determined through fluorescence-based immunohistochemistry (Fig. 3e). Hence, the location of SREBP1, SREBP2, SRB-1, and STAR was relatively tracked in HGSOC tissues of optimum, borderline, and high cohorts. Laser confocal micrographs of Fig. 3e unveiled the presence of SREBP1 in the cytoplasm as well as the nucleus of optimum and borderline HGSOC tissues. SREBP2 resided in the cytoplasm of optimal HGSOC tissues while an additional nuclear localization was noted among borderline HGSOC tissues. High HGSOC tissues displayed reduced SREBP1-expression within the cytoplasm in addition to delineating prominent nuclear SREBP2 localization (Fig. 3e). For SRB-1 proteins, sparse cytoplasmic appearances were noted among optimal HGSOC samples which became remarkably detectable among borderline tissues. On the other hand, membrane positive stains for SRB-1 were notable among high cohort tissues (Fig. 3e). Again, distinct cytoplasmic accumulation of STAR could be noticed in high cohort tissues which were missing in the borderline and optimal ones (Fig. 3e). Therefore, a function dependent translocation of SREBP2, SRB-1, and STAR was ascertained for the tissues of different cholesterol cohorts. Reconfirmation of this was further achieved by quantitative estimation of cohort-specific HMGCR activities. Graphical anecdote in Fig. 3f revealed the highest HMGCR activities among tissues of the high HGSOC cohort which also displayed nuclear SREBP2 as opposed to the borderline and optimal samples. This observation aligned with the established notion of SREBP2 activation being indispensable for the translation of the HMGCR enzyme. Moreover, SRB-1 enriched membranes in the HGSOC tissues of high cohort hinted at its function in enabling blood cholesterol uptake which lead to its increase within tissues. Therefore, increased cholesterol turnover among HGSOC patients of high cohort resulted from accelerated rates of *de novo* cholesterol synthesis as well as exogenous uptake. These results bridged the effects of deranged PI3K/Akt/mTOR signaling with abnormal cholesterol synthesis in HGSOC scenario thereby proposing their candidature as ‘screening markers’ for carboplatin therapy response assessment.

### Dysregulated PI3K/Akt/mTOR signatures are Directly Proportional to Compromised Carboplatin Response in HGSOC

Optimum (n=3), borderline (n=5), and high (n=10) HGSOC cohort tumors were primary cultured for addressing the cumulative impact of dysregulated PI3K/Akt/mTOR and cholesterol aberrance upon their carboplatin responses (Fig. 3a and S4a). As a requisite, MTT assay was performed following 24 hours treatment of the respective *ex vivo* cultures with logarithmic range (0.1-100µM) of carboplatin concentrations (Fig. 3b). Enumerated 50% growth inhibitory concentrations (IC_50_) of carboplatin varied in a cohort-dependent manner (optimum-1.59±0.31µM, borderline-2.78±0.72µM, high-5.23±0.704µM) along with their distinct cholesterol profiles and HMGCR activities (Fig. 3b, 1f, S1d, and 2h). Herein, 50% cell growth in high cohort *ex-vivo* cultures was inhibited by 1.89 fold (p<0.0001) and 3.3 fold (p<0.0001) higher carboplatin concentrations compared to borderline and optimum groups. Accordingly, cholesterol abundant high HGSOC tissues with escalated HMGCR activities were least carboplatin responsive. Instead, borderline tissues with relatively less cholesterol and corresponding HMGCR activities were observably less carboplatin responsive. Again, cholesterol deficient optimal HGSOC tissues with low HMGCR activities responded best to carboplatin (Fig. 3b and S4b). Therefore optimum, borderline, and high cohorts were classified as carboplatin ‘responder’, ‘intermediate responder’, and ‘non-responder’ respectively (Fig. S4b). These profiles portrayed a differential therapy response that is specific to cholesterol profiles of individual patients.

Relative alterations in cell specific expressions of PI3K/Akt/mTOR candidates were furthermore investigated among the HGSOC cohorts by flow cytometry after subjection to a ‘Median Effective Dose (MED)’ of 2µM carboplatin (Fig. 3c and S4c). Thereafter, modulations in expressions of cholesterol metabolism regulators (SREBP1, SREBP2, SRB-1, STAR) along with upstream (Akt/ pAkt^Thr308^, mTOR, pmTOR^Ser2448^, p70S6K, P38MAPK) and downstream (HIF-1α, COX2, VEGF) prosurvival effectors were comparatively evaluated with carboplatin’s presence or absence (Fig. 3d and S5). In a Heat map, these molecules were hierarchically clustered as ‘first priority’, ‘second priority’, and ‘third priority’ references based on equivalent abundances to predict HGSOC carboplatin responses (Fig. 3d and S5). Accordingly, pAkt^Thr308^, mTOR, p70S6K, P38MAPK, HIF-1α, COX2, VEGF, pmTOR^Ser2448^, SREBP2, and SRB-1 were individually annotated as ‘first priority’ references. Untreated high HGSOC tumors contained 42.87%-pAkt^Thr308^, 16.70%-mTOR, 13.47%-p70S6K, 11.47%-P38MAPK, 28.45%-HIF-1α, 26.45%-COX2, 22.80%-VEGF, 32.43%-pmTOR^Ser2448^, 30.40%-SREBP2 and 33.80%-SRB-1positive cells which increased considerably to 60.55%, 38.20%, 27.28%, 17.10%, 31%, 30.41%, 42.40%, 53.55%, 47%, and 41.02% upon carboplatin MED administration. Borderline HGSOC tumors harboring less number of these cells, with or without MED exposure were intermediate carboplatin responders in relation to non-responsive high-cohort. Again, these cells were sparsely noted among carboplatin responsive optimal HGSOC tumors which hardly increased in the presence of carboplatin. Correspondingly, their Mean Fluorescence Intensities (MFIs) corroborated with their cohort-specific cellular abundances both in presence and absence of carboplatin (Fig. 3e). Following close in the hierarchy were the ‘second priority’ markers namely STAR and Akt which majorly constituted borderline (15.67%, MFI-412; 14.45%, MFI-394) and high (20.74%, MFI-562; 30.44%, MFI-682) HGSOC TMEs unlike optimum (6.70%, MFI-182; 5.23%, MFI-151). Carboplatin MED treatment raised their numbers among borderline (STAR-12.3%, MFI-598; Akt-30.44%, MFI-427) and high (STAR-41.2%, MFI-689; Akt-40.10%, MFI-720) groups as opposed to optimum (Fig. 3d and 3e). Unrelated expression patterns of SREBP1 among optimum (-carboplatin: 52.14%, MFI-685; +carboplatin: 33.47%, MFI-544), borderline (-carboplatin: 3.66%, MFI-124; +carboplatin: 2.89%, MFI-116), and high (-carboplatin: 0.44%, MFI-67; +carboplatin: 0.20%, MFI-27) HGSOC cohorts distinguished it as an obscure ‘third priority’ reference (Fig.3d and 3e). Hence, HGSOC carboplatin responses appeared to be in close alliance with their molecular diversities (Fig. S5).

To re-ascertain if carboplatin-induced Akt phosphorylation underlined these dysregulations, pAkt^Thr308^ co-expressions with prosurvival (upstream/downstream) effectors and cholesterol metabolism regulators were determined (Fig. 3d). Here also, pAkt^Thr308^ co-expressions with Akt, pmTOR^Ser2448^, P38MAPK, COX2, and STAR were designated ‘first priority’. Heat map indications suggested high-HGSOC samples to be enriched with such double positive cells unlike optimum (Fig. 3d). In the absence of carboplatin, high-HGSOC samples harboured 17%-Akt, 33%-pmTOR^Ser2448^, 10%-P38MAPK, 43.5%-COX2, and 15%-STAR positive cells expressing pAkt^Thr308^. These cells became highly prominent (30%-Akt+pAkt^Thr308^, 61.3%-pmTOR^Ser2448^+pAkt^Thr308^, 20.70%-P38MAPK+pAkt^Thr308^, 50.14%-COX2+pAkt^Thr308^, 18.02%-STAR+pAkt^Thr308^) with carboplatin MED administration thereby establishing the necessity of Akt activation to denounce drug effects. Carboplatin exposure thereafter increased their corresponding MFIs as well (Fig. 3e). Contrarily, borderline samples presented lesser double positive multitudes with lower MFI ratios which further depreciated in optimum samples (Fig. 3d and 3e). Next up, the ‘second priority’ markers including pAkt^Thr308^ co-expressions with mTOR, SREBP2, SRB-1, HIF-1α, p70S6K, and VEGF were respectively noted among 33.6%, 27.6%, 21.2%, 19.8%, 7.6%, and 20.82% cells of carboplatin untreated high-HGSOC samples (Fig. 3d). On ensuing carboplatin MED, pAkt^Thr308^ got increasingly detected among 35.7%-SREBP2, 27.43%-SRB-1, 24.6%-HIF-1α, 10.4%-p70S6K, and 22.56%-VEGF bearing cells. Conversely, mTOR (unphosphorylated) could be detected only among 7.89%-pAkt^Thr308^ positive cells. Similarly, borderline cohorts constituted by relatively fewer double positive cells which underwent an increase during carboplatin triggers. Relative MFI ratios varied accordingly which reaffirmed that Akt activation was essential for mTOR, SREBP2, SRB-1, HIF-1α, p70S6K, and VEGF expressions among borderline HGSOC TME as well (Fig. 3e). This pattern went missing in optimum samples, both with or without carboplatin (Fig. 3d and 3e). Increasing expression of pAkt^Thr308^ and pmTOR^Ser2448^ in conjugation with diminishing profiles of Akt and mTOR indicated HGSOC’s adaptation to carboplatin via Akt phosphorylation (Fig. 3e). This overall molecular disharmony was further associated with lipid metabolism by validating PI3K/Akt/mTOR candidate expressions in context of SREBP1 (Fig. 3d and 3e). Formulated results evinced obscure SREBP1 co-expressions among optimum, borderline, and high HGSOC cohorts, with or without carboplatin challenge (Fig. 3d and 3e). Hence, SREBP1 co-expressions with all prosurvival effectors and cholesterol metabolism regulators were allotted ‘third priority’. This confirmed deregulated cholesterol metabolism as an intrinsic feature of all HGSOC cohorts which could possibly be an indicator of its carboplatin responsiveness.

These characteristic PI3K/Akt/mTOR derangements adhered with exhilaration of HMGCR activities and cholesterol titers particularly among the borderline and high HGSOC cohorts (Fig. 3f). High and borderline *ex vivo* cultures constitutively presented higher cholesterol levels (high-6.02 fold; borderline-2.24 fold) along with cumulatively raised HMGCR activities (high-4.24 fold; borderline-1.96 fold) unlike carboplatin responsive optimal cohort. Subjecting them to carboplatin subsequently accelerated their HMGCR activities by 6.4 and 2.74 folds; increasing cholesterol titers by 10.68 and 3.4 folds compared to optimum. Conclusively, a deregulated PI3K/Akt/mTOR signaling cascade revitalizes cholesterol enriched HGSOC tissues with poor carboplatin response.

### PI3K/Akt/mTOR mediated Cholesterol Sufficiency decides carboplatin-sensitivity among HGSOC cells

The cumulative effect of diminished cellular drug sensitivities and reduced chemoenhancement in compromising the carboplatin response of HGSOC cohorts was further scrutinized. Carboplatin MED treated *ex vivo* cultures were evaluated for drug sensitivities in terms of quantifying Pt-DNA adducts retention, DNA damage, ROS generation, mitochondrial(mt) membrane depolarization, and membrane cholesterol enrichment statuses (Fig. S6a). In flow cytometric enumeration of carboplatin-DNA adduct retaining potentials, 88.2% of optimal HGSOC cells were positive for Pt-DNA adducts in relation to borderline (~80.2%) and high (~78.4%) cohorts (Fig. 4a). Interestingly, a distinct 19.8% and 25.2% of Pt-DNA adduct negative cell populations were noted among borderline and high HGSOC cohorts (Fig. 4a). Persistence of such subcellular populations in the HGSOC TME correlated with their partial or complete carboplatin sensitivities. As Pt-DNA adducts formation damages DNA, comet assay was performed wherein carboplatin treated optimum *ex vivo* cultures espoused maximal drug induced DNA damage (Fig. 4b). Lesser extents of DNA damage as originally observed in borderline HGSOC cells visibly increased with carboplatin treatment but this was quantitatively less than optimal (Fig. 4b and S6b). Inherently greater DNA damage of high HGSOC cells remained unaltered with carboplatin shots; bestowing carboplatin insensitivity and unresponsiveness (Fig. 4b and S6b). As carboplatin also mediates oxidative stress to incur DNA damage, cohort-wise ROS levels were quantified by flow cytometry (Fig. 4c and S6b). Carboplatin’s presence made ROS levels reach the highest peaks in the optimum cohort, unlike borderline and high groups. Thus, a diminished histogram of optimal cells indicated carboplatin-triggered ROS evoked cell killing. Contrarily, elevated-basal ROS levels detected among borderline and high HGSOC tumors in the absence of carboplatin hardly escalated with drug treatment; justifying their diminished carboplatin responses (Fig. 4c and S6b). Since mt-membrane depolarization promotes ROS overproduction, JC-1 staining was performed to additively ascertain the same in flow cytometry (Fig. 4d and S6c). Pronounced mt-membrane depolarization was detected among optimum HGSOC cells with carboplatin encounter which was otherwise absent. Instead, the borderline cells emerged with relatively less depolarized membrane when carboplatin was ensued. Again, high HGSOC cells appeared to possess an already depolarized mt-membrane which barely changed with carboplatin treatment; hinting at decreased drug efficacy being the cellular adaptive measure to escape carboplatin cytotoxicity (Fig. 4d and S6c). Often, impeded drug entry across cholesterol enriched rigid membranes depletes intracellular carboplatin levels to aggravate this scenario. Thus, membrane rigidities of e*x vivo* cultured cells were gauged spectrofluorimetrically by LAURDAN staining, both prior to and after carboplatin MED administration (Fig.4e). A shift in laurdan fluorescence emissions from 500nm to 450nm among high HGSOC cells confirmed membrane rigidification which intensified upon carboplatin MED exposure. This was relatively toned down among untreated borderline HGSOC cells which increased nominally upon carboplatin treatment. Conversely, optimal HGSOC cells possessed fluid membranes as the LAURDAN fluorescence emissions peaked at 500nm which diminished subtly with carboplatin administration (Fig. 4e). Interestingly, membrane rigidification aligned with cholesterol enrichment patterns of optimum, borderline, and high HGSOC cohorts. Therefore, cholesterol abundant high HGSOC cells bearing rigid membranes were identified as carboplatin non-responder or carboplatin insensitive. Borderline HGSOC cells with moderate cholesterol levels and less rigid membranes surfaced as intermediate carboplatin responders or ‘partially carboplatin sensitive’. Subsequently, optimum HGSOC tumors with depreciating cholesterol turnovers and fluid membranes were carboplatin responders or carboplatin sensitive. Thus, reprogrammed cholesterol metabolism stemming from the deregulated PI3K/Akt/mTOR pathway primordially determines the carboplatin sensitivity and response of HGSOC tumors which could be exploited for designing pre-therapy screening rationales.

**Fig. 4:**
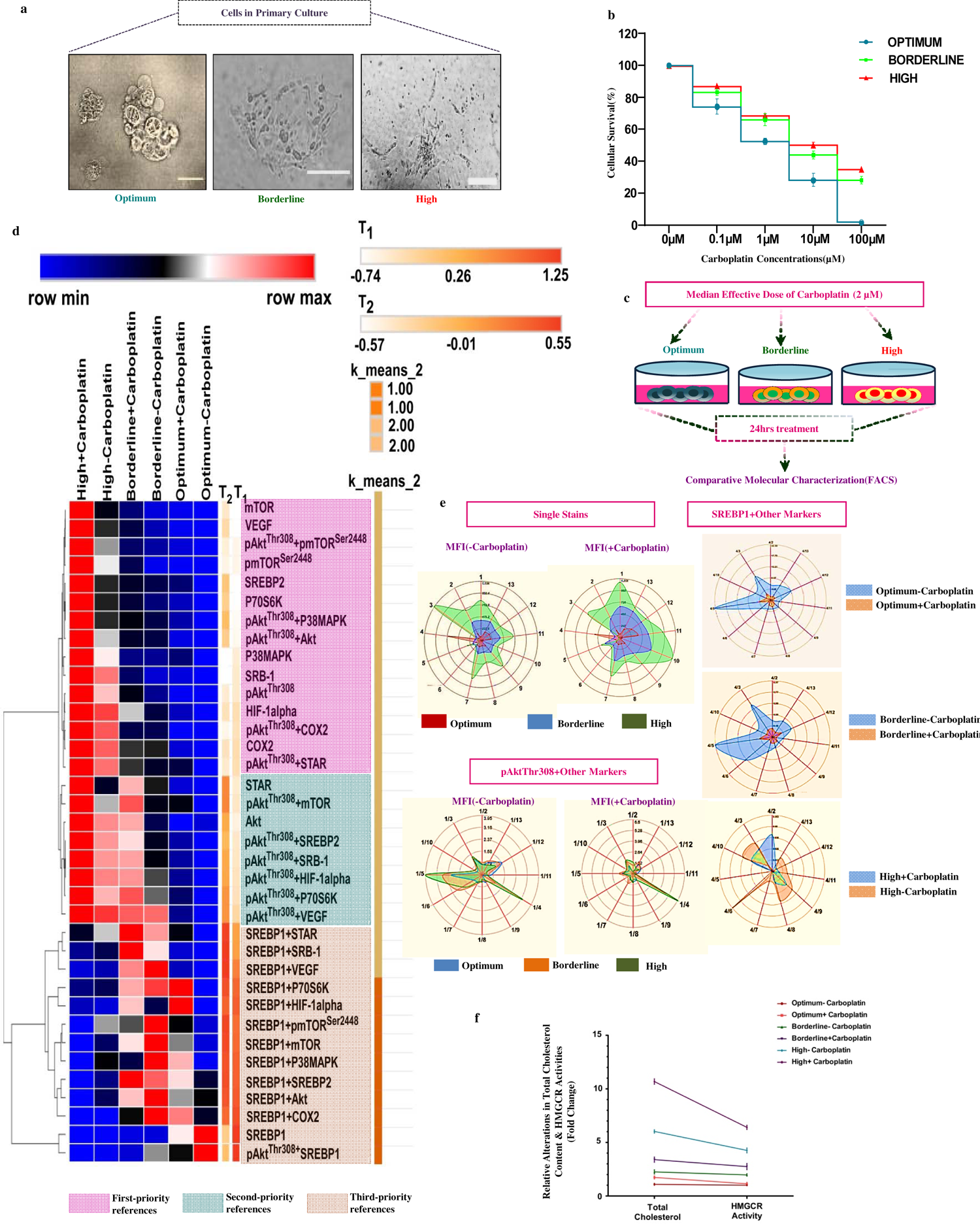
Deregulated PI3K/Akt/mTOR status along with enhanced cholesterol level in HGSOC is representative of poor carboplatin response: (a) Micrographic anecdotes of *ex vivo* cultured cells from tissues of optimum, borderline and high HGSOC cohorts. (b) Cohort wise variance of carboplatin response represented graphically. (c) Outline of the plan of work for flow cytometry study. (d) Heat Map representing relative abundance of cells positively expressing the selected PI3K/Akt/mTOR candidates including cholesterol metabolism regulators, singly as well as doubly. (e) Radar Charts depicting the corresponding Mean Fluorescence Intensities (MFI) for single stains (1-pAkt^Thr308^, 2-Akt, 3-mTOR, 4-SREBP1, 5-p70S6K, 6-P38MAPK, 7-HIF-1α, 8-VEGF, 9-COX2, 10-pmTOR^Ser2448^, 11-SREBP2, 12-SRB-1, 13-STAR) and MFI-ratios for double stains cohort specifically. (f) Graphical representations of relative alterations in tissue cholesterol levels and HMGCR activity profiles for the HGSOC cohorts. All results are represented as Mean±S.D.

**Fig. 5:**
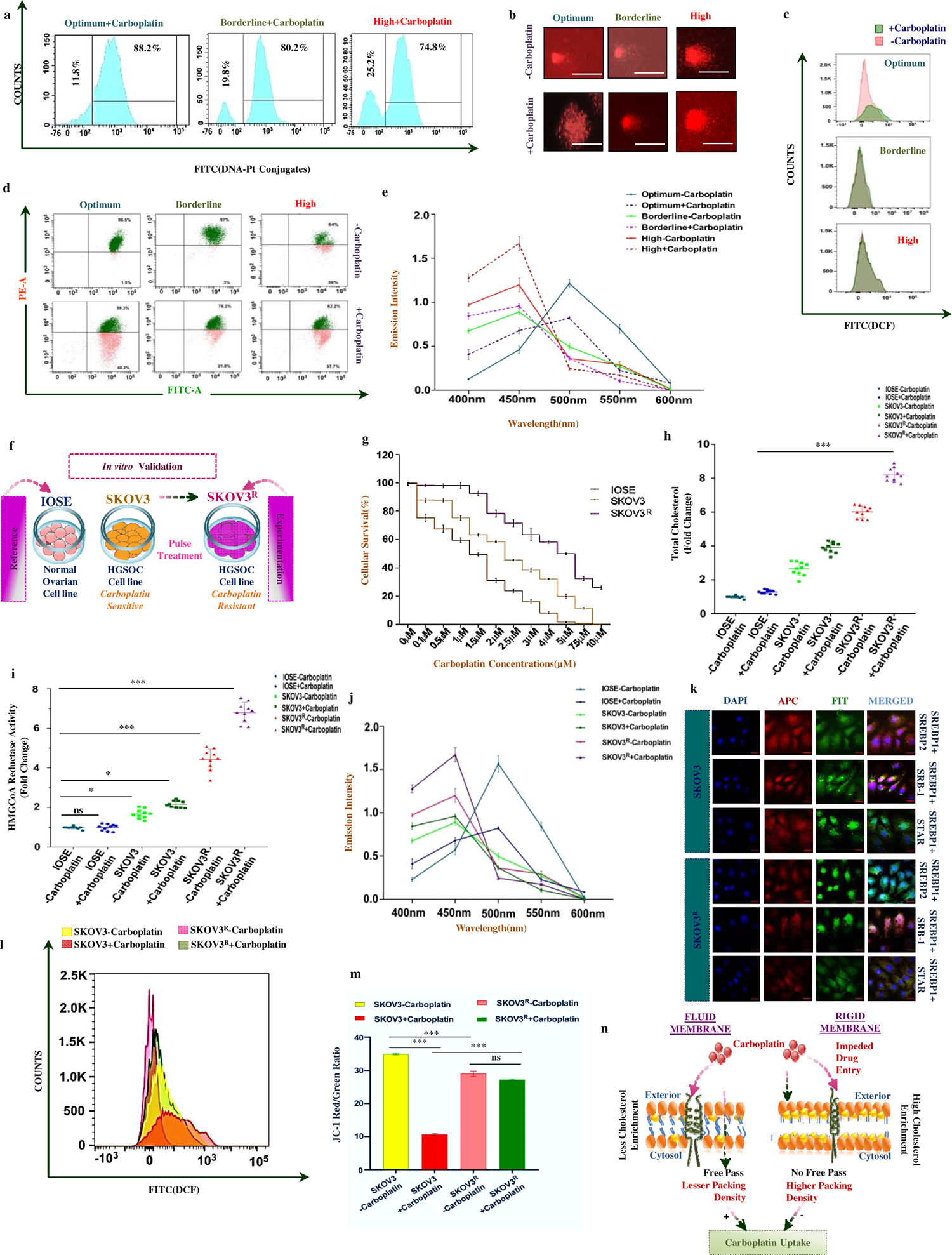
Poor carboplatin response correlates with high cholesterol profiles: (a) Cohort-specific Pt-DNA adducts retention. (b) Comet Assay micrographs depicting DNA damage among HGSOC cohorts. (c) Cohort-wise ROS generation with or without carboplatin treatment. (d) JC-1 membrane potential variance with drug treatment among HGSOC cohorts. (e) Fluorescence intensity peaks representing membrane rigidification among HGSOC cohorts. (f) *In vitro* plan of work. (g) MTT results of IOSE, SKOV3 and SKOV3^R^. Relative cholesterol levels among the *in vitro* models. (i) Trends of HMGCR activity profiles of IOSE, SKOV3 and SKOV3^R^. (j) Relative membrane rigidities of *in vitro* cell lines with and without carboplatin treatment. (k) Tissue specific location of SREBP2, SRB-1 and STAR among SKOV3 and SKOV3^R^ cells. (l) Carboplatin induced ROS generation among SKOV3 and SKOV3^R^. (m) JC-1 red to green ratio representing membrane depolarization SKOV3 and SKOV3^R^. (n) Summary of study. All results were represented as Mean ± S.D with *p<0.05, **p=0.05 and ***p<0.0001.

Ahead of concluding the study, we verified these endpoints in an *in vitro* setup wherein a HGSOC cell line SKOV3 was used besides a normal cell line, IOSE as a reference. SKOV3^R^, a 2.1 fold carboplatin-resistant subline of SKOV3 was developed by ‘pulse treatment’ (Fig. 4f). Besides enumerating IC_50_ values, the carboplatin response of SKOV3 and SKOV3^R^ was ascertained in the form of membrane rigidification, ROS generation, and mt-membrane depolarization. As per MTT assay results, SKOV3^R^ with an IC_50_ of 6µM was carboplatin-resistant whereas SKOV3 and IOSE cells with IC_50_ values of 3.1µM and 1.7µM were mildly carboplatin-resistant and carboplatin sensitive respectively (Fig. 4g). SKOV3^R^ furthermore recorded higher intracellular cholesterol levels and HMGCR activities compared to SKOV3 and IOSE along with maximal rigidities in the membrane (Fig. 4h, 4i, and 4j). With carboplatin treatment, these parameters increased manifold among SKOV3^R^ cells, unlike SKOV3 and IOSE cells. These findings closely corroborated with carboplatin-triggered greater ROS titers and raised mt-membrane depolarization rates of SKOV3^R^ in contrast to SKOV3 (Fig. 4l). SKOV3^R^ evinced nuclear SREBP2, membrane SRB-1, and cytoplasmic STAR proteins unlike SKOV3 cells as revealed from immunofluorescence results (Fig. 4k). Carboplatin responsive SKOV3 was ‘drug sensitive’ while SKOV3^R^ with minimal carboplatin response was ‘drug resistant’. Higher cellular cholesterol rigidified membranes to hinder carboplatin entry and accumulation; compromising drug response to promote resistance as schematically depicted in Fig. 4n. Cumulatively, these results established that reprogrammed cholesterol metabolism and PI3K/Akt/mTOR deregulations synergistically enable HGSOC tumors to escape carboplatin cytotoxicity thereby paving a way for therapy-failure.

## DISCUSSION

HGSOC therapy with carboplatin, either singly or in combination with paclitaxel or primary debulking surgeries is a ‘one track’ approach for clinicians. Limited treatment options and lack of technological amenities to predict the probable outcomes of conventional therapy make disease management cumbersome for clinicians [10–13]. Such an emergency demands the establishment of reliable pre-therapy screening strategies for which a holistic understanding of the disease is a prima facie. We, therefore, attempted to characterize the HGSOC tumors with key focuses on their intrinsic attributes-PI3K/Akt/mTOR signaling, cholesterol metabolism, and platinum therapy response. Aggressive HGSOC cells reinforce their upregulated PI3K/Akt/mTOR signaling for accelerating *de novo* cholesterol synthesis and exogenous uptake to increase their sterol reliance for proliferating autonomously [36]. To ascertain its relevance in our patient scenario, we initiated this study with an attempt to find out if abysmal cholesterol biosynthesis is an exclusive issue for HGSOCs. For this purpose, we formulated comparative blood cholesterol profiles for HGSOC patients in reference to normal volunteers and other gynaecological cancer patients. Accordingly, all the study subjects could be distributed among ‘optimum’, ‘borderline’, and ‘high’ cholesterol cohorts. Herein, HGSOC candidates came up with significant disparities, especially within the borderline and high cohorts. They presented poorly differentiated tumors with aggressive pathology and relatively high tissue cholesterol content as well unlike optimum cohort subjects. Our results detected high cohort patients with raised blood and tissue cholesterol levels and poorly differentiated aggressive tumors. Our findings definitively enabled the identification of aberrant blood cholesterol levels as an intrinsic feature of HGSOCs. In fact, a collateral surge in tissue cholesterol content vis-à-vis loss in tissue differentiation further strengthened this notion. With a sample size of 31 patients, our findings could not sufficiently ascertain a risk range for ovarian tissue cholesterol that could be referential for its aggressive nature. In light of blood cholesterol, we determined that in HGSOC patients the titer may unusually shoot up from a basal lower limit of 165 mg/dl to an upper limit of about 301mg/dl which was absent in other cancer patients or normal non-cancer volunteers. In a nutshell, abnormal blood cholesterol levels were identified as a hallmark of the HGSOC scenario.

It is well known that aggravated malignancies absolve therapy stress by their upregulated oncogenic signaling and concomitantly reprogrammed metabolism [23–25]. Thus, we proceeded with comprehending the molecular underpinnings of an aggressive HGSOC TME that led to abnormal cholesterol production which was also reflected in blood. In this regard, the TCGA database reporting HGSOC cases with mutated PI3K/Akt/mTOR candidates directly or indirectly regulating cholesterol metabolism justified the study rationale. This laid the foundation for selective scrutiny of Akt/pAkt^Thr308^, mTOR/pmTOR^Ser2448^, p70S6K, P38MAPK, HIF-1α, COX2, VEGF, SREBP1, SREBP2, SRB-1, and STAR protein in a cohort specific manner. With raised Akt, mTOR, p70S6K, P38MAPK, HIF-1α, COX2, and VEGF proteins, high-HGSOC cohort espoused PI3K/Akt/mTOR mediated cholesterol metabolism reprograming in the effect of its high tissue cholesterol profiles despite indistinct pAkt^Thr308^ and pmTOR^Ser2448^ expressions. Again, SREBP2, the prime decider of cholesterol turnovers in tissues came up as an exclusive growth essential factor for HGSOCs owing to its indispensability for patient survival. We discerned an interaction of SREBP2 with mTOR, p70S6K, HIF-1α, SRB-1, and STAR in these samples. This further confirmed that the upstream or downstream prosurvival effectors along with cholesterol metabolism regulators orchestrated PI3K/Akt/mTOR signaling to promote cholesterol biosynthesis and uptake. Furthermore, function dependent localization of SREBP2, SRB-1, and STAR among high cohort HGSOC tissues suggested the same. Nuclear SREBP2 corroborated with the heightened HMGCR activity of high HGSOC cohort which corresponded with enhanced tissue cholesterol turnover. In this way, PI3K/Akt/mTOR dysregulation rewired cholesterol metabolism in high-HGSOC cohorts by driving more endogenous cholesterol biosynthesis. Moreover, high cohort tissues also delineated membrane SRB-1proteins which hinted at their tendency to harbour cholesterol exogenously. Heightened cytoplasmic STAR expressions in high cohorts affirmed increased cholesterol metabolism and the consequent hypoxia that might have induced higher HIF-1α expressions. The signaling myriad potentiated HGSOC aggressiveness because Co-IP and *in silico* studies also confirmed the same. These were moderately noted among borderline cohorts thereby appropriating deregulated cholesterol dynamics for aggressive HGSOCs only. Unaltered cohort-wise and diminished tissue specific expressions of SREBP1 reinstated that remodelled cholesterol profile in HGSOC TME originates from dysregulated PI3K/Akt/mTOR signaling.

Previously upregulated PI3K/Akt/mTOR was reported to diversify the HGSOC TME with cellular clones that respond differentially to carboplatin [29, 30]. Therefore, PI3K/Akt/mTOR effectors could be promising as ‘predictive markers’ of therapy outcomes. We ratified this in 2-D *ex vivo* primary cultures of HGSOC tissues collected from all three cohorts. Interesting observations were obtained in the presence as well as absence of carboplatin. High HGSOC tumors displayed carboplatin based selective increase of pAkt^Thr308^ along with Akt, pmTOR^Ser2448^, P38MAPK, COX2, and STAR in first-priorities. In second-priority, high cohorts also delineated similar survival advantages for cells co-expressing pAkt^Thr308^ alongside mTOR, SREBP2, SRB-1, HIF-1α, p70S6K, and VEGF with carboplatin shots. These tumors with the highest IC_50_ accustomed themselves to carboplatin encounter via Akt and mTOR phosphorylation. High cohort tumors were carboplatin non-responsive and were overpopulated with cells positively expressing pAkt^Thr308^, mTOR, p70S6K, P38MAPK, HIF-1α, COX2, VEGF, pmTOR^Ser2448^, SREBP2, SRB-1, STAR, and Akt. These HGSOC cells survived effortlessly despite the carboplatin challenge, probably by means of ‘Darwinian Selection’. With feeble Pt-DNA adducts retention, high cohort HGSOC cells had no intensification of DNA damage, ROS levels, and mt-membrane depolarization on carboplatin treatment. Persistence of such subcellular cohorts catered to partial or complete carboplatin insensitivities whereas the optimum cohort with the lowest IC_50_ lacked these anecdotes and was carboplatin-responsive/ sensitive.

This pilot study is observational involving protein studies only. However, the end-point observations may aid in strategizing therapy with novel ‘screening regimes’ as well as ‘drug-repurposing’ procedures. Earlier studies have reported a generalized association of upregulated PI3K/Akt/mTOR and platinum resistance in malignancies [25–36]. Our study connects these aspects with cholesterol metabolism in the HGSOC scenario because carboplatin evoked HMGCR activities to heighten cholesterol biosynthesis among the high HGSOC cohort. Herein, raised cholesterol levels possibly favoured carboplatin insensitivity by rigidifying cell membranes to impede drug entry. As evident from LAURDAN staining, most rigid membranes were found in high cohort HGSOC as opposed to borderline and optimum cohorts. These results corroborated in an *in vitro* setup; confirming the *modus operandi* of therapy escape. The unmet need for screening tumors prior to therapy could therefore be addressed by nominal investigation of HGSOC biopsy samples for their protein expressions of PI3K/Akt/mTOR candidates besides quantification of their cholesterol contents. Additionally, the evaluation of cholesterol reference range in the blood can be cost-effectively translated to clinics for segregating responders from non-responders before prescribing carboplatin. This would also aid in preventing unwanted dose exposure among patients thereby curtailing mortalities caused by greater ‘off-target’ effects in chemotherapy.

Thus, our inferences successfully proposed and verified the credibility of deregulated PI3K/Akt/mTOR effectors and reprogrammed cholesterol profiles as ‘screening markers’ for predicting the carboplatin response of HGSOC tumors.

## CRediT authorship contribution statement

**Elizabeth Mahapatra:** Conceptualization, data curation, formal analysis, investigation, methodology, software, validation, visualization, writing original draft. **Arka Saha:** Data curation, formal analysis, investigation, methodology, software, visualization. **Niraj Nag:** Methodology, resource, software, visualization. **Animesh Gope:** Software, formal analysis. **Debanjan Thakur**: Data curation. **Manisha Vernekar:** Conceptualization, resource. **Jayanta Chakrabarti:** Conceptualization, resource. **Mukta Basu:** Methodology, software, validation. **Amit Pal:** Resource. **Sanghamitra Sengupta:** Conceptualization. **Sutapa Mukherjee:** Conceptualization, data curation, fund acquisition, project administration, resources, supervision, validation, writing-reviewing & editing.

## DATA AVAILABILITY

All data is available upon request from the corresponding author.

## ACKNOWLEDGMENTS

We are grateful to Dr. Sib Sankar Roy, Chief Scientist & Head, Cell Biology & Physiology Division, CSIR-IICB, Kolkata, INDIA for his kind help in providing SKOV3 cell line. We would also extend vote of thanks to Mr. Subhabrata Guha, Junior Research Scholar, Dept. of Signal Transduction and Biogenic Amines, CNCI, Kolkata for his help in constructing overlay curves. We heartily thank Ms. Shalini Upadhayay for her cooperation in recording and analysing all the flow cytometry results portrayed in this manuscript.

## Abbreviations

HMGCR: 3-Hydroxy-3-methylglutaryl-coenzyme A Reductase
Ca-125: Cancer antigen-125
Chl: Cholesterol
COX2: Cyclooxygenase 2
HGSOCs: High-Grade Serous Ovarian Cancers
HIF-1α: Hypoxia Inducible Factor-1α
mTOR: mammalian Target of Rapamycin
mt-membrane: Mitochondrial membrane
non-neoadjuvant chemotherapy: Non-NACT
P38MAPK: P38MAPKinase
P70S6K1: P70 Ribosomal protein S6 kinase
PI3K: Phosphoinositide3 Kinase
Akt: Protein Kinase B
SRB1: Scavenger Receptor Class B Type 1
STAR: Steroidogenic Acute Regulatory
SREBP2: Sterol Response Element Binding Protein 2
TME: tumor microenvironment
VEGF: Vascular Endothelial Growth Factor

## Supplementary Figure Legends

**Fig. S1: Aberrant Blood and Tissue Cholesterol Profiles are intrinsic to HGSOC:** (a) Pictorial description of the Rationale and work plan followed for the results depicted in Figure 1. (b) Relative fold change in the total plasma cholesterol levels among optimum, borderline and high cohorts of normal, other cancer and HGSOC subjects. (c) Noteworthy anecdotes of HGSOC histopathologies from optimum, borderline and high cohorts respectively. (d) Relative fold change in the total tissue cholesterol levels among optimum, borderline and high cohorts of other cancer and HGSOC subjects. Results of (b) and (d) are represented as Mean ± S.D with *p<0.05, **p=0.05 and ***p<0.0001.

**Fig. S2: HGSOC tissues with enhanced Cholesterol turnover displayed upregulated PI3K/Akt/mTOR candidates:** (a) Distribution of most frequently mutated genes among the 28 HGSOC cases reported in TCGA Database. (b) Relative Band Intensities of pmTOR, mTOR, p70S6K, pAkt, Akt and P38MAPK normalized against β-actin. (c) Relative Band Intensities of VEGF, HIF-1α and COX2 normalized against β-actin. (d) Relative Band Intensities of SREBP1, SRB-1, SREBP2 and STAR normalized against β-actin. Results were represented as Mean ± S.D (*p<0.05, ***p<0.0001).

**Fig. S3: Increased Cholesterol Levels results from exacerbated HMGCR activities promoted by activation of Cholesterol Metabolism Regulators in HGSOC:** (a) Correlation Scatter plots depicting importance of co-expression of SREBP2 with SRB-1, STAR, and SREBP1. (b) Survival dependency of non serous patients vs serous patients upon SREBP2. (c) A plausible interactome of PI3K/Akt/mTOR candidates regulating cholesterol metabolism effectors to reprogram its metabolism in HGSOC.

**Fig. S4: Deregulated PI3K/Akt/mTOR status along with enhanced cholesterol level in HGSOC is representative of poor carboplatin response:** (a) A Schematic outline of primary culture protocol. (b) Cohort specific carboplatin response status in relation to their IC_50_ values. (c) Gating Patterns followed for the flow cytometry data analysis.

**Fig. S5: Variance of marker abundance in relation to IC_50_ values of HGSOC cohorts:** trend curves representing the variance of ic_50_ values for each HGSOC cohorts with their relative abundance (% positive cells) of first-priority, second-priority and third-priority markers. Each replicates have been plotted individually.

**Fig. S6: Poor carboplatin response correlates with high cholesterol profiles:** (a) Study design represented schematically. (b) Relative fold change of Comet Tail length among HGSOC cohorts in presence or absence of carboplatin. (c) Comparative Profile of ROS levels among HGSOC cohorts. (d) JC-1 Red to Green ratio depicting membrane depolarization of HGSOC cohorts with carboplatin treatment. Gating patterns of (e) Pt-DNA adduct retention study (f) ROS quantification and (g) JC-1 staining by flow cytometry. (h) Carboplatin sensitivities of HGSOC cohorts in relation to their respective IC_50._ All results were represented as Mean ± S.D with *p<0.05, **p=0.05 and ***p<0.0001.

