## Supplementary Figures for "Dysregulated Cholesterol Metabolism with Anomalous PI3K/Akt/mTOR pathway Predicts Poor Carboplatin Response in High Grade Serous Ovarian Cancer"

### Slide 1
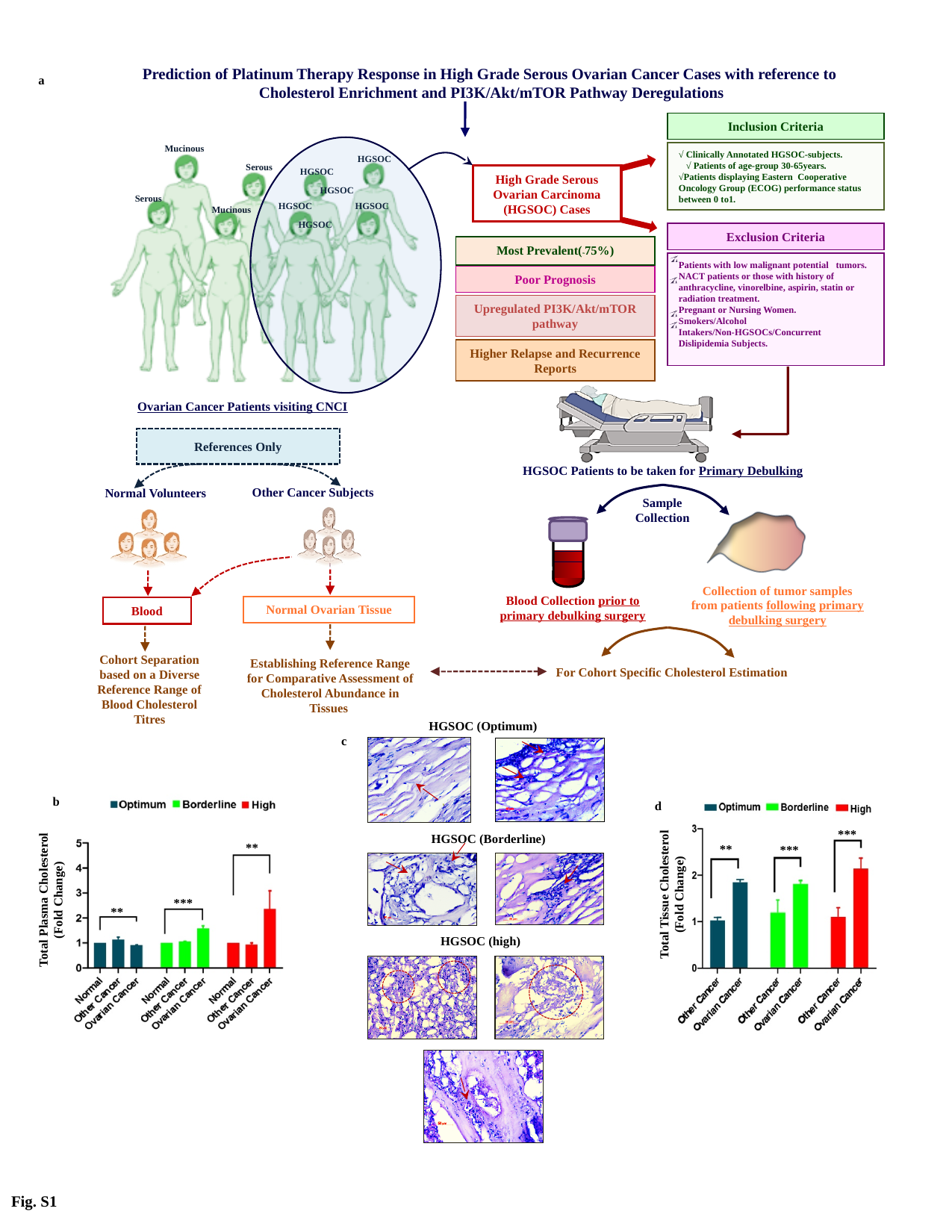

Prediction of Platinum Therapy Response in High Grade Serous Ovarian Cancer Cases with reference to Cholesterol Enrichment and PI3K/Akt/mTOR Pathway Deregulations
Inclusion Criteria
√ Clinically Annotated HGSOC-subjects.
 √ Patients of age-group 30-65years.
√Patients displaying Eastern Cooperative Oncology Group (ECOG) performance status between 0 to1.
Exclusion Criteria
Patients with low malignant potential tumors.
NACT patients or those with history of anthracycline, vinorelbine, aspirin, statin or radiation treatment.
Pregnant or Nursing Women.
Smokers/Alcohol Intakers/Non-HGSOCs/Concurrent Dislipidemia Subjects.
High Grade Serous Ovarian Carcinoma (HGSOC) Cases
Most Prevalent(_˜_75%)
Poor Prognosis
Upregulated PI3K/Akt/mTOR pathway
Higher Relapse and Recurrence Reports
Ovarian Cancer Patients visiting CNCI
Mucinous
HGSOC
Serous
HGSOC
HGSOC
Serous
HGSOC
HGSOC
Mucinous
HGSOC
References Only
HGSOC Patients to be taken for Primary Debulking
Other Cancer Subjects
Normal Volunteers
Sample Collection
Collection of tumor samples from patients following primary debulking surgery
Blood Collection prior to primary debulking surgery
Normal Ovarian Tissue
Blood
Cohort Separation based on a Diverse Reference Range of Blood Cholesterol Titres
Establishing Reference Range for Comparative Assessment of Cholesterol Abundance in Tissues
For Cohort Specific Cholesterol Estimation
a
HGSOC (Optimum)
c
HGSOC (Borderline)
HGSOC (high)
**
***
**
Total Plasma Cholesterol
(Fold Change)
b
Total Tissue Cholesterol
(Fold Change)
***
**
***
d
Fig. S1

### Slide 2
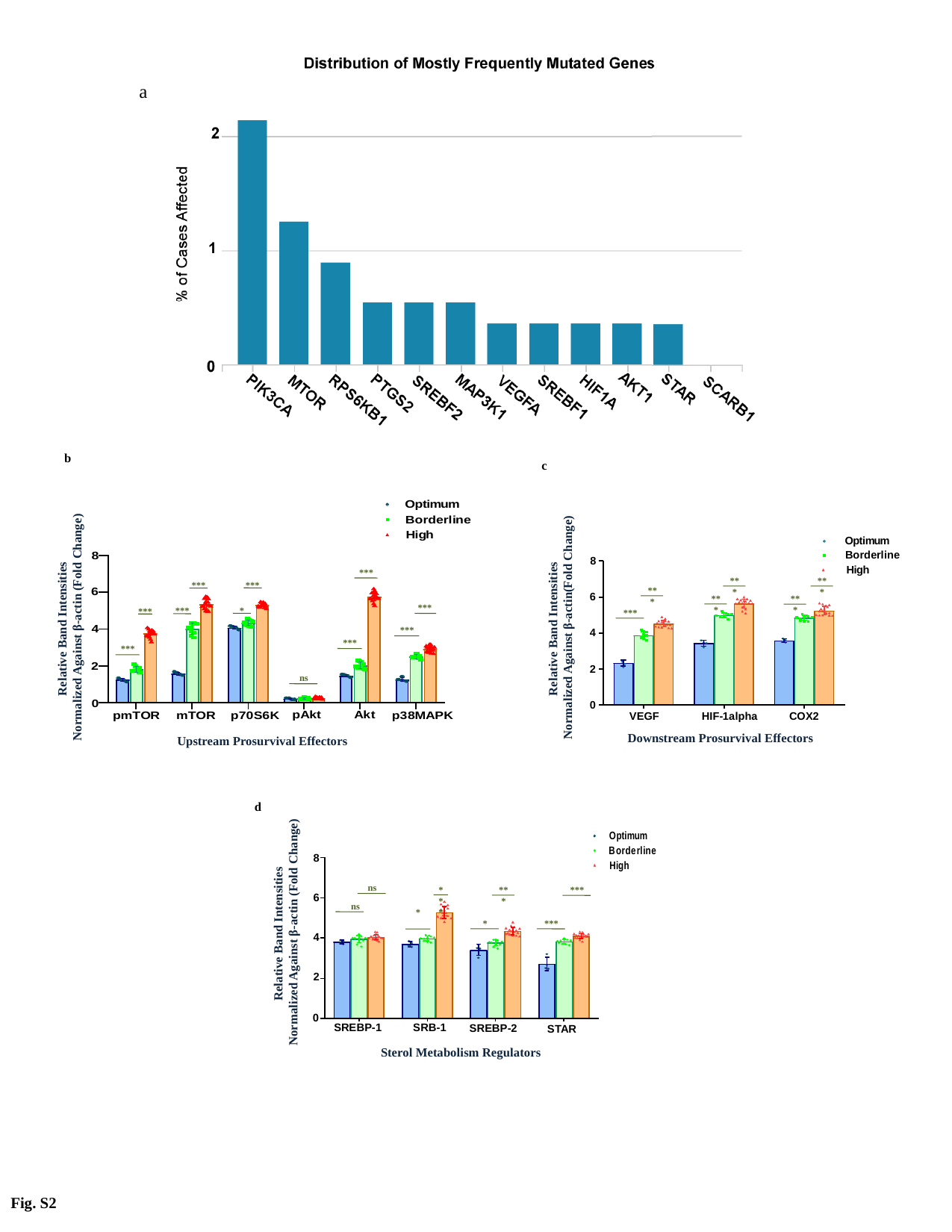

a
b
c
Relative Band Intensities
Normalized Against β-actin (Fold Change)
Upstream Prosurvival Effectors
***
***
***
***
***
*
***
***
ns
***
***
Relative Band Intensities
Normalized Against β-actin(Fold Change)
Downstream Prosurvival Effectors
***
***
***
***
***
***
d
Relative Band Intensities
Normalized Against β-actin (Fold Change)
Sterol Metabolism Regulators
ns
ns
***
*
***
*
***
***
Fig. S2

### Slide 3
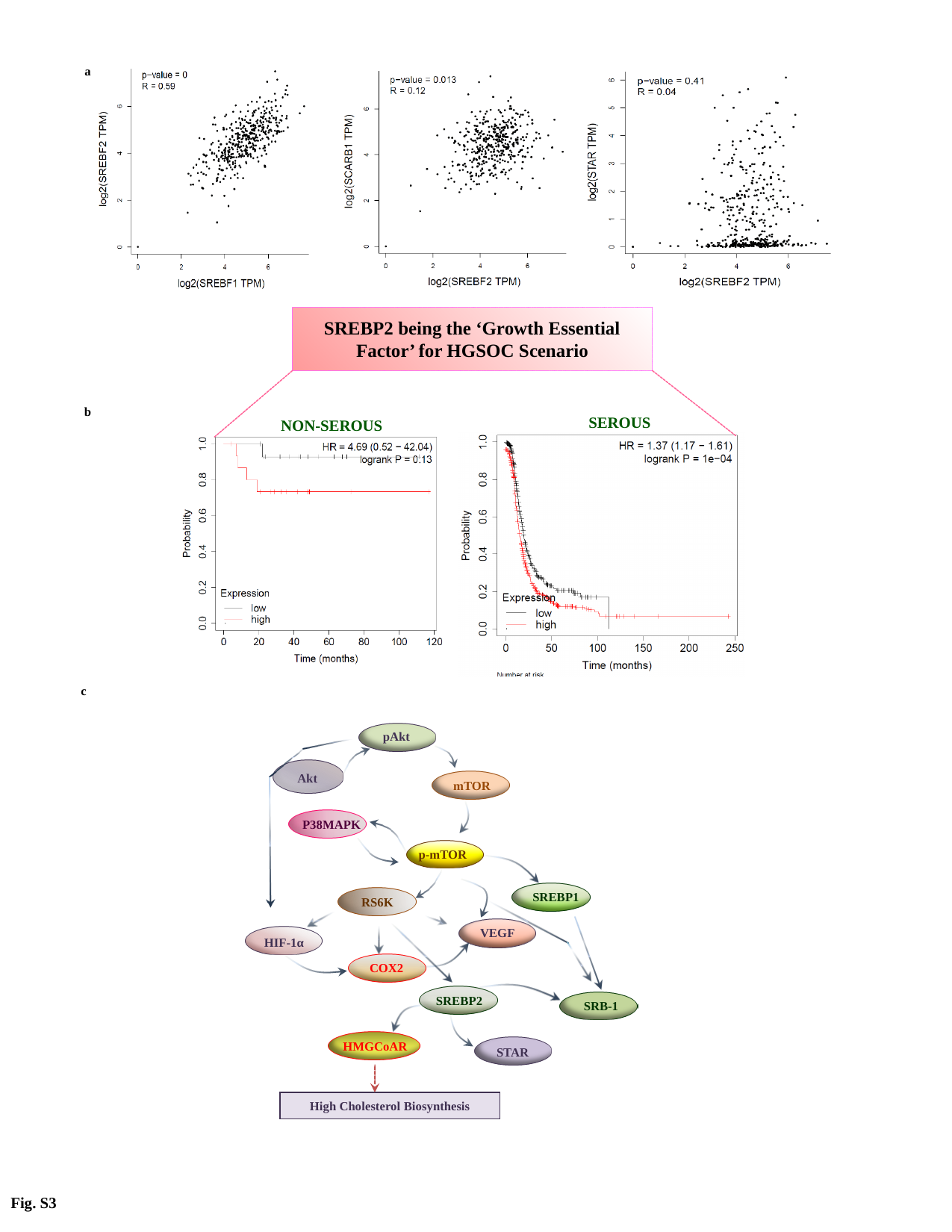

a
SREBP2 being the ‘Growth Essential Factor’ for HGSOC Scenario
b
SEROUS
NON-SEROUS
c
pAkt
Akt
mTOR
P38MAPK
p-mTOR
SREBP1
RS6K
VEGF
HIF-1α
COX2
SREBP2
SRB-1
HMGCoAR
STAR
High Cholesterol Biosynthesis
Fig. S3

### Slide 4
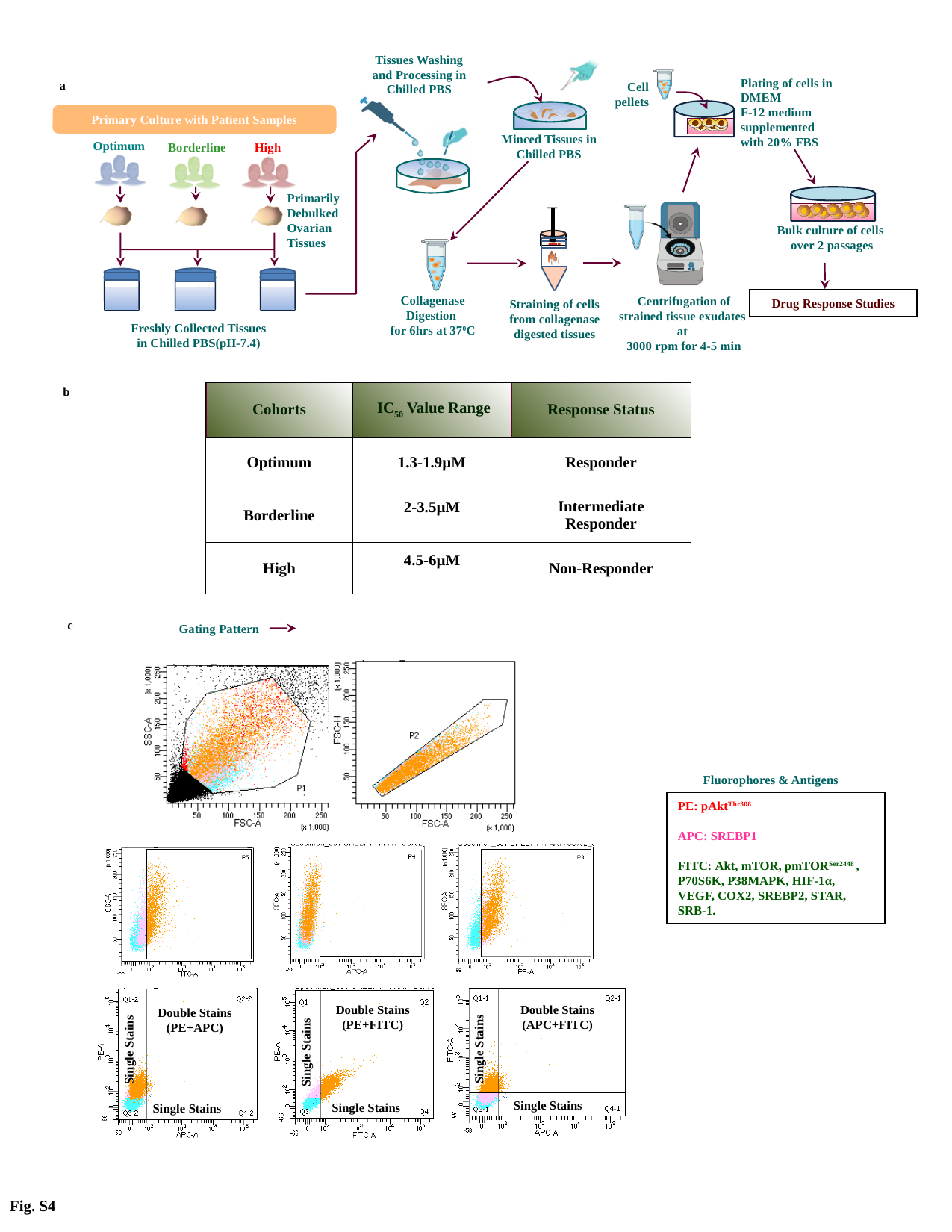

Tissues Washing and Processing in Chilled PBS
Minced Tissues in Chilled PBS
Cell
pellets
Plating of cells in DMEM
F-12 medium supplemented with 20% FBS
Primary Culture with Patient Samples
Optimum
Borderline
High
Primarily
Debulked
Ovarian
Tissues
Centrifugation of strained tissue exudates at
3000 rpm for 4-5 min
Straining of cells from collagenase digested tissues
Bulk culture of cells
over 2 passages
Collagenase Digestion
for 6hrs at 37^0^C
Drug Response Studies
Freshly Collected Tissues
in Chilled PBS(pH-7.4)
a
b
| Cohorts | IC_50_ Value Range | Response Status |
| --- | --- | --- |
| Optimum | 1.3-1.9µM | Responder |
| Borderline | 2-3.5µM | Intermediate Responder |
| High | 4.5-6µM | Non-Responder |
c
Gating Pattern
Double Stains
(PE+FITC)
Double Stains
(APC+FITC)
Double Stains
(PE+APC)
Single Stains
Single Stains
Single Stains
Single Stains
Single Stains
Single Stains
Fluorophores & Antigens
PE: pAkt^Thr308^
APC: SREBP1
FITC: Akt, mTOR, pmTOR^Ser2448^ , P70S6K, P38MAPK, HIF-1α, VEGF, COX2, SREBP2, STAR, SRB-1.
Fig. S4

### Slide 5
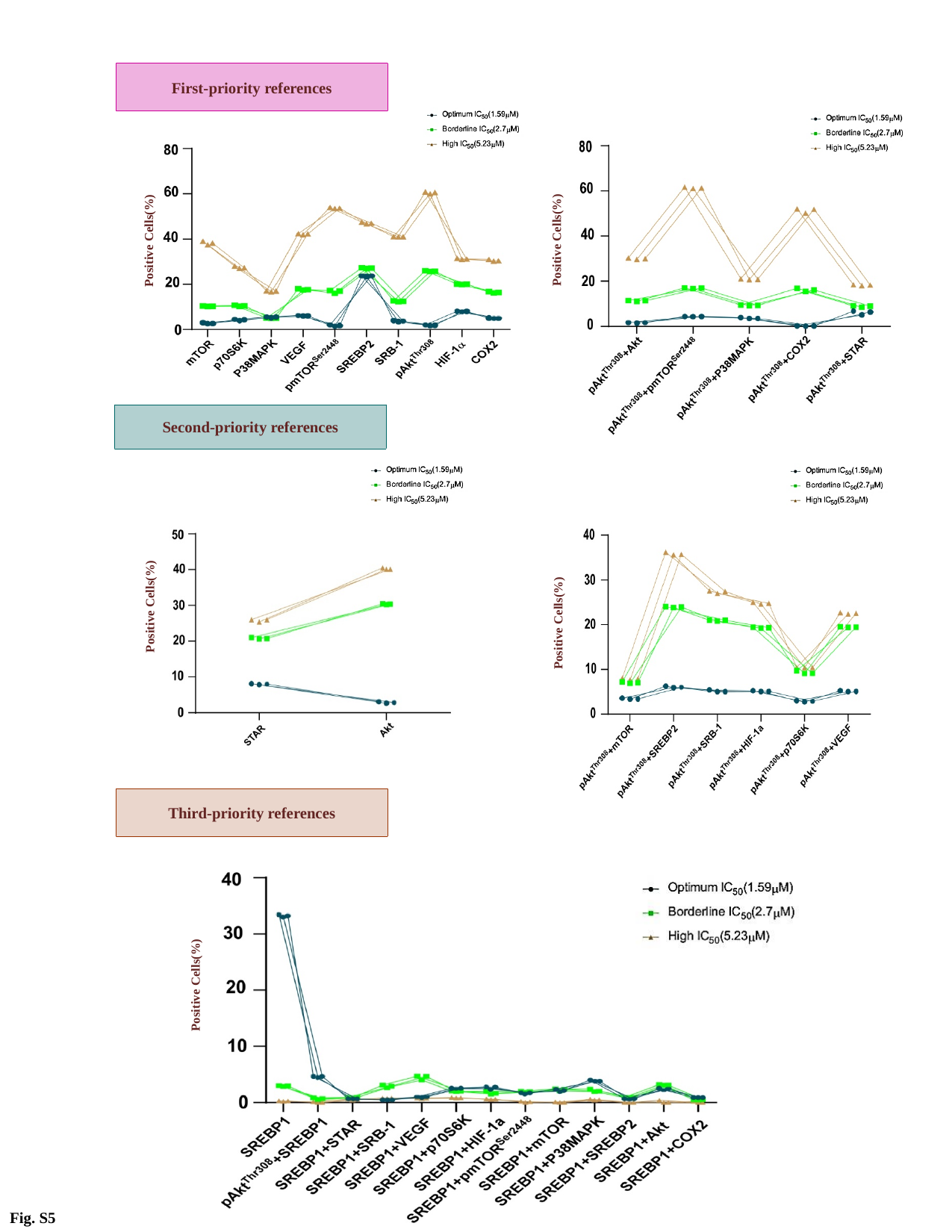

First-priority references
Positive Cells(%)
Positive Cells(%)
Second-priority references
Positive Cells(%)
Positive Cells(%)
Third-priority references
Positive Cells(%)
Fig. S5

### Slide 6
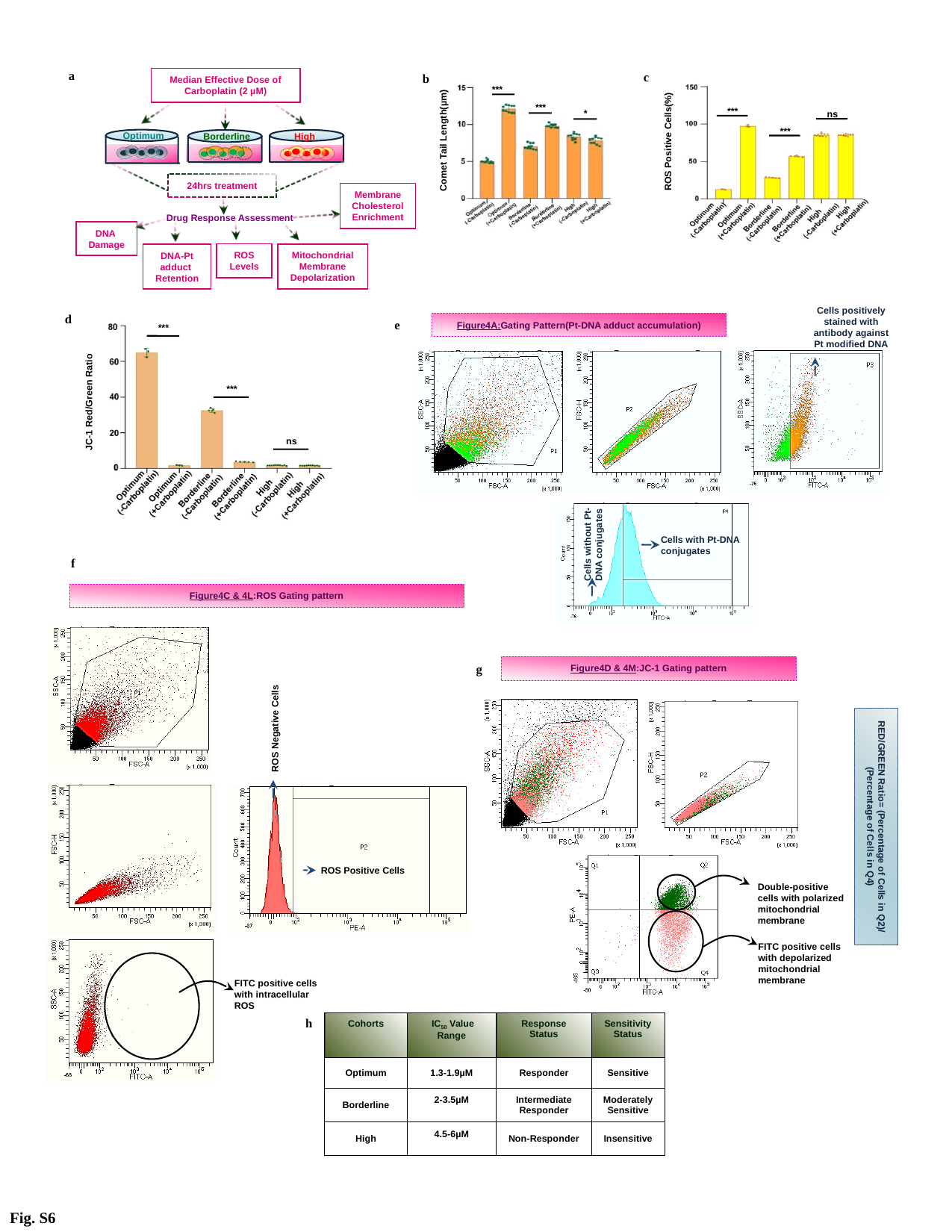

a
Median Effective Dose of Carboplatin (2 µM)
Optimum
High
Borderline
24hrs treatment
Drug Response Assessment
Membrane Cholesterol Enrichment
DNA Damage
ROS
Levels
Mitochondrial Membrane Depolarization
DNA-Pt adduct
Retention
c
***
ns
***
ROS Positive Cells(%)
b
Comet Tail Length(µm)
***
***
*
Cells positively stained with antibody against Pt modified DNA
Cells without Pt-DNA conjugates
Cells with Pt-DNA conjugates
Figure4A:Gating Pattern(Pt-DNA adduct accumulation)
e
d
***
JC-1 Red/Green Ratio
***
ns
f
Figure4C & 4L:ROS Gating pattern
ROS Negative Cells
ROS Positive Cells
FITC positive cells with intracellular ROS
Figure4D & 4M:JC-1 Gating pattern
g
Double-positive cells with polarized mitochondrial membrane
FITC positive cells with depolarized mitochondrial membrane
RED/GREEN Ratio= (Percentage of Cells in Q2)/ (Percentage of Cells in Q4)
h
| Cohorts | IC_50_ Value Range | Response Status | Sensitivity Status |
| --- | --- | --- | --- |
| Optimum | 1.3-1.9µM | Responder | Sensitive |
| Borderline | 2-3.5µM | Intermediate Responder | Moderately Sensitive |
| High | 4.5-6µM | Non-Responder | Insensitive |
Fig. S6
